# Modeling the acutely injured brain environment *in vitro*

**DOI:** 10.1101/2025.01.08.631921

**Authors:** Dulce María Arzate, Víctor Álvaro, Ana Victoria Prádanos-Senén, Lucía Gallego, Alba Méndez Alejandre, Diana Manzano, Celia Llorente-Sáez, Rocío Bartolomé, Alessia Ferrarini, Juan Antonio López, Elisa Murenu, Benedikt Berninger, Felipe Ortega, Gabriel Piedrafita Fernández, Magdalena Götz, Christophe Heinrich, Sergio Gascón

## Abstract

A major challenge to study the regenerative potential of the injured brain is the limited access to this organ in vivo. To address this, we developed an innovative gliosis model that, although established *in vitro*, originates from a genuine injury *in vivo*. The model relies on reactive glia acquiring enhanced adhesion, facilitating their rapid adaptation to *in vitro* conditions, where it faithfully recapitulates key features of brain injury. These include a secretome associated with injury pathways, degenerative responses like neuronal death and neuroinflammation, and regenerative processes such as progenitor proliferation, recruitment and commitment to oligodendrocytes. Moreover, the exposure of adult glial cells to this culture medium recapitulates their acquisition of multipotency observed in both mouse and human injured brains. Finally, our approach allows studying glia-to-neuron reprogramming, a process challenging to tackle *in vivo*. Consequently, we present a novel tool for exploring stem cell dynamics and regenerative behaviors in CNS pathology.

## INTRODUCTION

Acute brain injuries are major causes of mortality and neurological disability worldwide ^1^. These conditions compromise the blood-brain barrier (BBB), disrupting blood flow and leading to oxygen deprivation ^2^, which severely affects neurons due to their strong dependence on mitochondria to sustain their metabolic functions ^3^. Furthermore, BBB disruption causes the leakage of molecules and immune cells from the bloodstream into the brain ^4^, with the former initiating a gliosis response and the latter amplifying the pro-inflammatory environment ^5^. This, along with reactive oxygen species (ROS) generated by the subsequent restoration of vascular flow and mitochondrial activity, contributes to neuronal death ^6^.

Despite these adverse effects, the primary goal of neuroinflammation is beneficial ^7^, aiming to remove pathogens and debris throughout activated astrocytes and myeloid cells. In addition, certain cytokines and growth factors released by immune cells can stimulate adult neural stem cells’ (aNSCs) proliferation and differentiation into astrocytes, neurons, or oligodendrocytes ^8,9^, suggesting a certain degree of neuro-regenerative capacity. Furthermore, in zebrafish, inflammatory signals are needed for brain regeneration after lesion ^10^. Also consistent with this notion, cumulative evidence indicates that reprogramming glial cells into induced neurons (iNs) is more efficient in injured than in intact brains ^11,12^. However, unlike organs such as the skin, where it can promote repair ^13^, inflammation in the mammalian CNS more frequently results in tissue damage and neuronal death, as previously discussed.

To effectively promote CNS regeneration over degeneration, it is essential to understand the divergent inflammatory responses discussed above, where aNSCs, glial cells, macrophages, and neurons all play important roles. A major challenge here lies in the limited access to observe and manipulate the injured brain *in vivo*, including the extracellular compartment where the reactive cells reside. To overcome these limitations, researchers have turned to simpler models of reactive cultures *in vitro*, typically mono-lineage, such as microglia activated by LPS, or astrocytes rendered reactive by exposure to cytokine-rich media ^14,15^ or TGF-β ^16^, among other stimuli. In some cases, these models comprise multiple cell types, particularly astrocytes, microglia, and neurons ^17^, to better reflect the complexity of CNS disease processes. Still, while these approaches offer highly valuable insights, they provide a limited representation of the cellular diversity and intricate interactions present in the injured brain. On the other hand, organotypic cultures ^18^, which retain more of the original cell and tissue architecture, are complex to maintain and manipulate *ex vivo*.

To address these limitations, we established a novel *in vitro* model of gliosis guided by two key principles: 1) gliosis is initiated *in vivo*, with reactive glial cells subsequently cultured under controlled *in vitro* conditions, and 2) cell selection exploits the markedly enhanced adhesive properties exhibited by all the reactive cellular lineages in the injured brain, enabling their exceptionally rapid attachment. As a result, the cultures are inherently reactive and comprise the cell types typically found in the lesioned brain, including activated microglia, reactive astrocytes, oligodendrocytes, oligodendrocyte precursor cells (OPCs), pericytes and immune cells from the bloodstream. Furthermore, the cultures reproduce a complex molecular landscape of proteins, metabolites, and signaling pathways characteristic of the injured brain. A similar approach to the one described in this study has previously been used to elucidated the role of galectins in promoting multipotency in adult reactive astrocytes following injury ^19^ (see also ^20,21^) and demonstrated the capacity of SEZ-derived progenitors to migrate into the injured motor cortex and transition into neurosphere-forming cortical astrocytes ^22^. Here, we expand on our understanding of this multicellular reactive culture (RC), while providing a detailed description of the method to enable others to apply it in their own research.

## RESULTS

### Establishing a multicellular reactive culture that represents the primary glial lineages characteristic of the injured brain

Our study aimed to model the gliotic environment *in vivo* as faithfully as possible through an *in vitro* approach. For the lesion we used a modification of the stab wound injury model in mice ^12^, selected for its simplicity and effectiveness in inducing gliosis and disrupting both the BBB and the compartmentalization of brain tissue, which are key features of acute lesions ^23^. As detailed in the Materials and Methods section and illustrated in Figure 1A and Supplementary Video 1, sixteen short and precise incisions were made on the somatosensory-visual cortex area.

**Figure 1.**
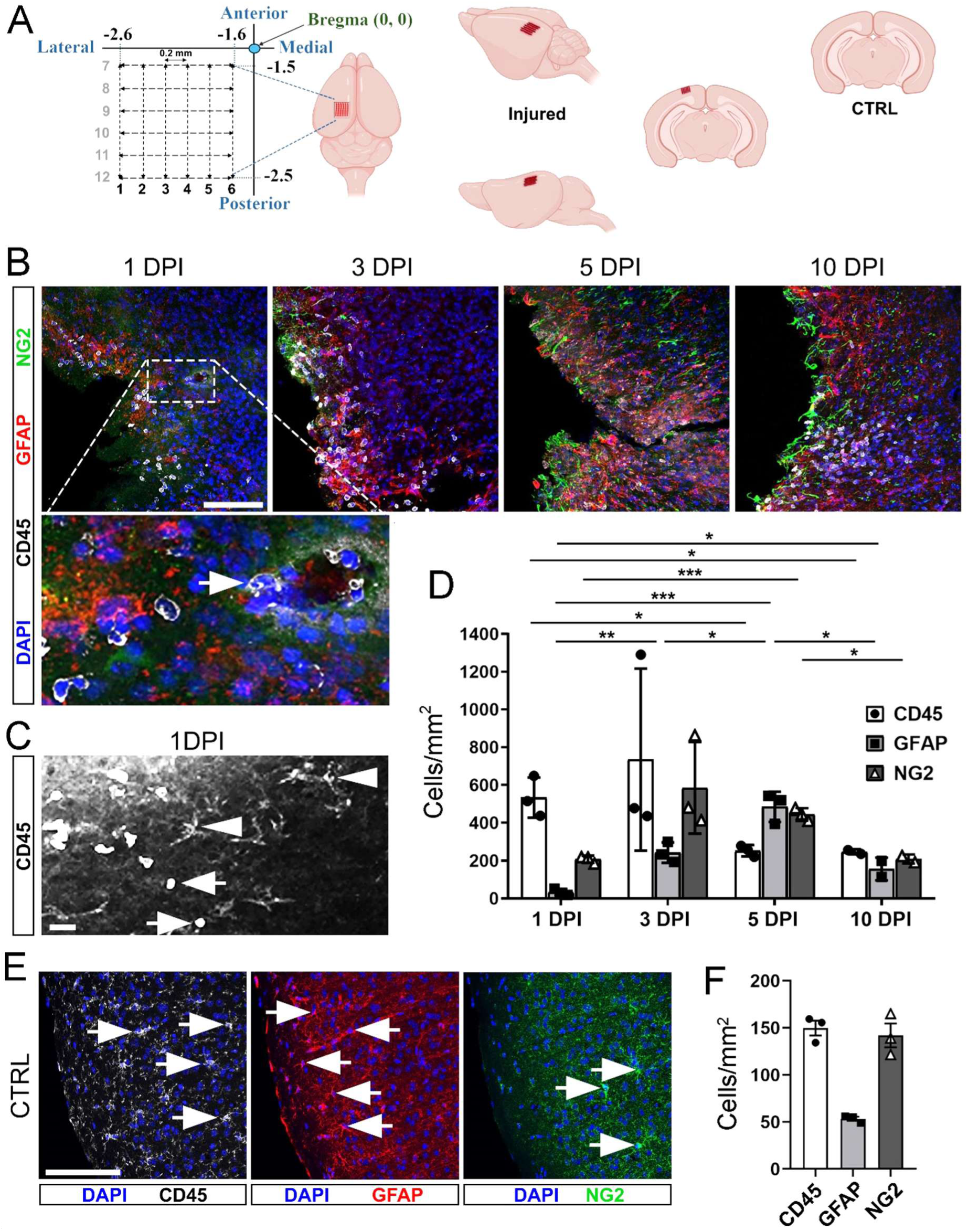
Stab wound injury model used to initiate a reactive glia culture. **(A)** Schematic representation of the stab wound lesion in the mouse adult cortex. The lesion was induced with a lancet positioned at Bregma coordinates (anteroposterior −1.6 mm to −2.5 mm, mediolateral ±1.5 mm to ±2.5 mm, and dorsoventral 0.5 mm). The lancet was moved ∼1 mm back and forth along the anteroposterior axis to create six traces, followed by six additional tracks along the mediolateral axis (12 total; see also Supplementary Movie 1). **(B)** Micrographs of the injured cortex at different DPI illustrate the presence of the previously mentioned glial lineages. Arrow in inset shows a CD45⁺ immune cell extravasating from a blood vessel. **(C**) CD45^+^ myeloid cells comprise rounded blood-derived monocytes with high CD45 expression (CD45-high, depicted by arrows) and ramified resident microglia with lower CD45 expression (CD45-low, depicted by arrowheads). **(D**) Quantification of immunoreactive cells per mm² across the 1–10 DPI conditions. **(E)** Immunostaining of an uninjured control (CTRL) brain slice showing myeloid cells (CD45^+^; white), reactive astrocytes (GFAP^+^; red), and OPCs (NG2^+^; green), along with nuclei (DAPI; blue). Arrows depict examples of cells from specific lineages. **(F**) Quantification of immunoreactive cells per mm² in CTRL brains. Statistical analysis: One-way ANOVA followed by a post-hoc Tukey test. Statistically significant differences are indicated as follows: * p<0.05; ** p<0.01; *** p<0.001. Scale bars: 100 µm in B and D; 50 µm in E.

The incisions were shallow (0.5 mm) to prevent damage to the white matter, and across the anteroposterior and mediolateral axes to cover an entire area of 1 square millimeter. After completing the procedure, the bone was replaced to aid healing. We then monitored the presence of distinct glial cell populations at 1, 3, 5, and 10 days post-injury (DPI) using immunohistochemistry (Figure 1B, C).

The number of myeloid CD45^+^ cells rapidly increased within 24 hours post-lesion, peaked at day 3, and declined by day 5 (Figure 1B-D). These included both rounded cells with high expression of CD45 (hereafter referred to as CD45-high) identified as blood-derived immune cells ^24^, and polygonal, ramified resident microglia with lower CD45 levels (referred to as CD45-low; see Figure 1C, where arrows and arrowheads distinguish monocytes from microglia). Some CD45-high cells were found breaching blood vessels walls (Figure 1B inset). Concurrently, the numbers of GFAP^+^ reactive astrocytes and NG2^+^ cells, likely OPCs, increased by 3 DPI and remained elevated until 5 DPI (Figure 1B, D). By day 10, cell proportions balanced (Figure 1B, D) and returned to levels similar to uninjured brains (Figure 1E, F). Thus, *in vivo* enrichment times for cultures are likely 1-5 DPI for myeloid cells and 3-5 DPI for NG2^+^ OPCs and GFAP^+^ reactive astrocytes. However, this will depend on the temporal progression of the glial lineages *in vitro*.

We subsequently established RCs from lesions at the time points mentioned above (Figure 2A). For this purpose, mice were euthanized at each designated interval, and their brains extracted. Lesion areas were then isolated by axial cuts, producing ∼3 mm thick coronal slices containing the damaged tissue (Figure 2A, middle panel), distinguishable by its typically brighter appearance. Then we subdivided each slice into two thinner coronal sections, containing the lesion, and next resected the damaged tissue for culturing, avoiding collecting white matter and meninges. After that, we processed and digested the tissue as per Materials and Methods, and plated the cells on poly-D-lysine (PDL)-coated glass covers in 24-well-plates at one hemisphere per well. Injured cortex cells readily adhered to coverslips within 2 hours, while cells from non-lesioned controls or from contralateral cortices showed minimal or no attachment (Figure 2A, right panel).

**Figure 2.**
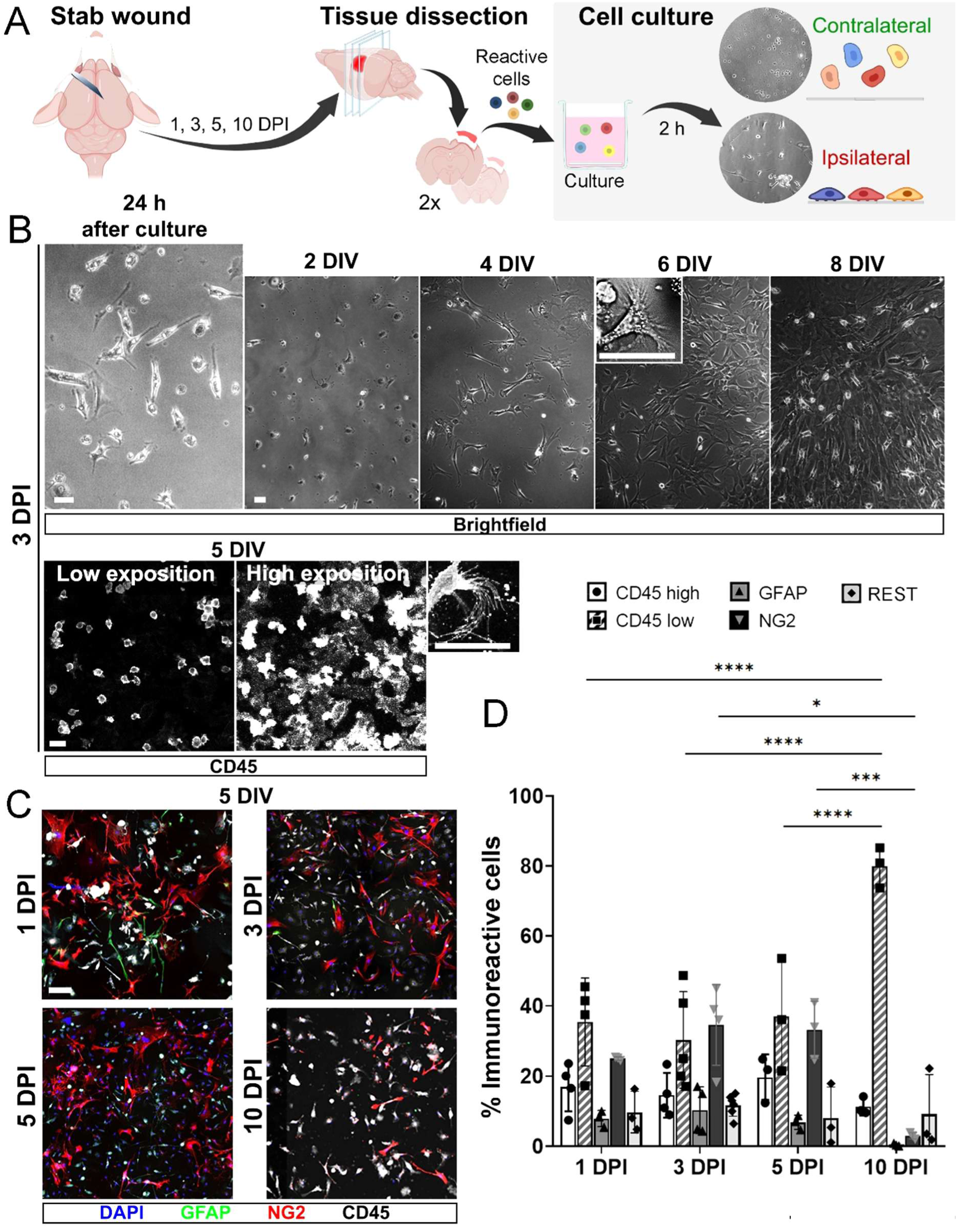
Reactive cultures established at early post-lesion time points contain the reactive glial populations observed *in vivo* after brain injury. (**A**) Schematic representation of the protocol for establishing RCs from the adult mouse injured cortex at 1, 3, 5, or 10 DPI. Brightfield images on the right show cells from injured tissue adhering within two hours, whereas those from contralateral samples do not. (**B**) Brightfield images in upper panels show the temporal progression of a 3 DPI reactive culture over 1, 2, 4, 6, and 8 DIV. Lower panels display CD45 immunostaining at low and high exposure, with rounded cells showing high CD45 levels and amoeboid/branched microglia showing low CD45. Insets in the 6 DIV brightfield and 5 DIV CD45 staining panels highlight filopodia-bearing microglia. (**C**) Mosaic images (merged with LAS X software) of CD45 (white), GFAP (red), and NG2 (green) immunoreactive cells in RCs from 1, 3, 5, and 10 DPI, examined after 6 DIV. (**D**) Quantification of markers, distinguishing CD45-high from CD45-low cells as a percentage of DAPI-positive cells. Cells not identified by any of the markers are labeled as REST. Statistical analysis: One-way ANOVA followed by a post-hoc Tukey test. Significant differences: *p<0.05; **p<0.01; ***p<0.001; **p<0.0001. Scale bars: 50 µm.

Micrographs from brightfield microscopy at different time points revealed rapid progression of cultures across all tested post-lesion intervals, reaching confluence by 8 days *in vitro* (DIV; see examples of 3 DPI cultures in Figure 2B, upper panels). By 4-6 DIV, distinct cell morphologies were observed, including microglia-like ameboid shapes with filopodia (inset in upper panels of Figure 2B-6DIV). The presence of myeloid cells was confirmed by immunocytochemistry, identifying CD45-high and CD45-low populations (Figure 2B, lower panels), likely corresponding to infiltrating immune cells and resident microglia, respectively ^24^, with the latter frequently exhibiting filopodia (Figure 2B, inset in lower panels). To further confirm these lineages, we used knock-in mice expressing RFP under the Ccr2 promoter (Figure S1), which is highly expressed in blood-derived monocytes and macrophages but in low levels in microglia ^25^. In RCs derived from these mice, CD45-high cells consistently showed strong immunoreactivity to RFP and IBA1 (Figure S1, micrographs). Since the concurrent expression of CD45, Ccr2, and IBA1 at high levels is associated with infiltrating macrophages in the brain ^25,26^, this observation further supports the blood origin of CD45-high cells.

Subsequent analyses of RCs from 1 to 5 DPI, and maintained for 5 DIV, revealed an approximate 3:2:1 ratio of CD45, NG2⁺ and ⁺, GFAP⁺ cells (Figure 2C, D), closely mirroring *in vivo* proportions at 1 and 3 DPI (Figure 1D). A shift towards an increased number of NG2⁺ cells was observed at 5 DPI, with microglia predominating at 10 DPI (Figure 2C, D). Therefore, RCs between 1 and 3 DPI exhibit the greatest cellular diversity and may represent the most suitable stage for future applications. In this regard, our previous study ^22^ showed that some migrating SEZ-derived progenitors could reach the motor cortex by 3 DPI, and be collected in the cultures. This appears unlikely under the current method, given the long distant from the SEZ to the stab wound area, located in the posterior somatosensory-visual. However, this possibility cannot be entirely ruled out at later post-injury time points.

### Cultures established one or three days post-injury contain a highly diverse population of glial cells

Consistent with previous observations, RCs established from 1 and 3 DPI exhibited the greatest glial diversity, prompting us to examine their cell composition in greater detail. To this end, we monitored RCs derived from lesioned tissues at 1DPI and 3DPI and cultured for 4, 8, and 20 days (Figure 3 and Figure S2). We observed an increase in DAPI^+^ nuclei over time, indicating active cellular proliferation (Figure S2A, B). Furthermore, the ratio of CD45^+^, NG2^+^, and GFAP^+^ cells remained steady around 3:2:1 at 4-5 DIV (see previous Figure 2C, D) and 8 DIV (see Figure 3A-C, G for 1 DPI, Figure 3D-F for 3 DPI, and G), regardless of when the cultures were established post-injury. Together, both observations indicate that cell proliferation occurs in the three glial lineages at both 1 and 3 DPI. Moreover, all the three glial populations continued to thrive through 20 DIV in all tested conditions (Figure 3C, right panel; Figure 3E, lower panel; Figure 3H and Figure S2C).

**Figure 3.**
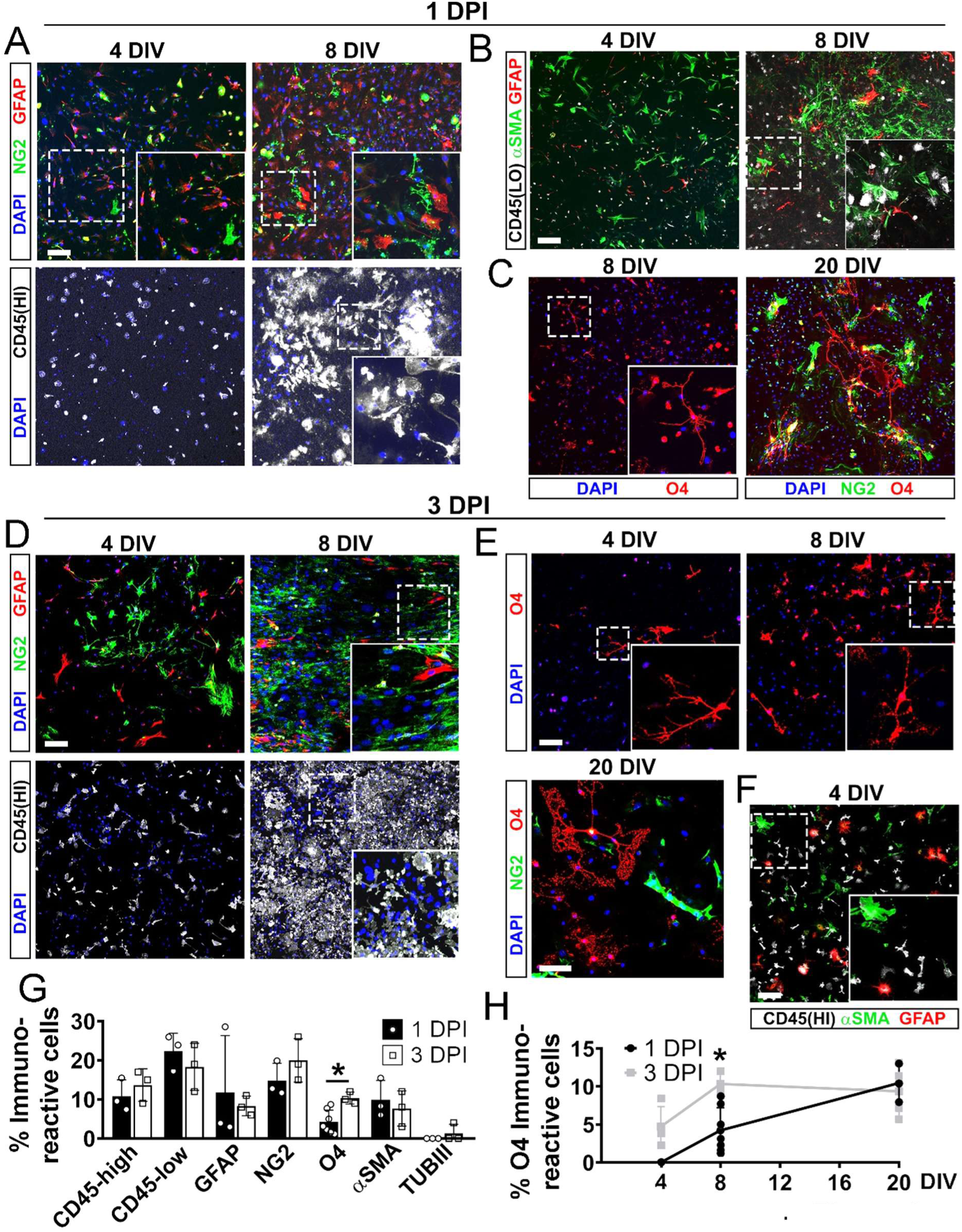
Reactive cultures established at 1 or 3 DPI exhibit extensive glial diversity. (**A**) Micrographs of NG2^+^ (green), GFAP^+^ (red) and CD45^+^ (white) cells at 4 DIV (left) and 8 DIV (right) in RCs established at 1 DPI, with magnified insets highlighting morphological details. Note CD45-high and CD45-low cells in the high-exposure (HI) CD45 immunostaining inset. (**B**) CD45 (white, LO: low exposure), α-SMA (green), and GFAP (red) immunostaining at 4 and 8 DIV in the same cultures. (**C**) O4^+^ cells at 8 DIV and both O4^+^ and NG2^+^ cells at 20 DIV in 1 DPI cultures, with inset showing a characteristic branched oligodendrocyte morphology. (**D**) NG2 (green), GFAP (red), and CD45 (white, high exposure) immunostaining in 3 DPI cultures at 4 DIV (left) and 8 DIV (right). (**E**) Immunostaining of the same cultures showing O4 (red) at 4 DIV (upper left) and 8 DIV (upper right), and both O4 (red) and NG2 (green) at 20 DIV (bottom). Note that at 20 DIV, oligodendrocytes exhibit complex morphology and lack NG2 expression, (**F**) Micrograph and inset showing CD45^+^ (white, high exposure), α-SMA^+^ (green), and GFAP^+^ (red) cells in 3 DPI cultures at 4 DIV. (**G**) Histogram of immunoreactive cell percentages for each marker in 1 DPI (black) and 3 DPI (gray) RCs. (**H**) Line graph showing earlier oligodendrocyte appearance in 3 DPI cultures. Data in G and H are mean ± SD (n = 3). Statistical analysis: One-way ANOVA followed by a post-hoc Tukey test, *p < 0.05. Scale bars: 75 μm.

Next, we further delved into the astroglial populations by assessing the expression of the pan-astrocyte marker S100β and found that approximately half of S100β^+^ cells did not express GFAP (Figure S3A). This suggests that the RCs harbor about twice as many astrocytes as detected by GFAP staining, exhibiting diverse reactivity profiles. Further highlighting this diversity, we identified astrocytes with varied morphologies (Figure S3B) and a rare but detectable subset co-expressing GFAP and NG2 (Figure S3C), previously reported in mouse ischemic brains ^27^. We then turned our attention to the CD45⁺ myeloid cells, and aimed to determine whether they were in a reactive state, as suggested by their expression of IBA1 (Figure S1). In fact, we found widespread expression of CD68 and CD11b (Figure S3D), being both markers of activated microglia and macrophages ^28^.

Beyond immune cells and astrocytes, our analyses also identified αSMA-immunoreactive cells (Figure 3B, F, G), likely representing pericytes derived from blood vessels ^29^, and O4-immunoreactive cells displaying branched morphologies (Figure 3C, E, G, H), presumably oligodendrocytes. Pericytes were detectable as early as 4 DIV and constituted approximately 10% of the cell culture by 8 DIV, regardless of whether the RCs originated from 1 or 3 DPI injured cortices (Figure 3B, F, and G). In contrast, the emergence of oligodendrocytes varied significantly with the timing of culture initiation. In cultures established at 1 DPI, O4^+^ oligodendrocytes were detectable by 8 DIV (∼4%) and abundant at 20 DIV (∼10%; Figure 3C, G, H), but entirely absent at 4 DIV (Figure 3H), thus suggesting they were not collected from the resected tissue but differentiated *in vitro*, likely from the cultured NG2^+^ OPCs.

Conversely, in RCs derived from lesions at 3 DPI, O4^+^ cells were detectable as soon as 4 DIV (∼5%; Figure 3E, H), suggesting that oligodendroglial differentiation began earlier, probably even *in vivo* prior to culture establishment. Notably, by 20 DIV, oligodendrocytes in RCs exhibited extensive branching, a hallmark of maturation (Figure E, lower panel), indicating that RCs support a full oligodendrogenesis program, as observed within the acute inflammatory environment *in vivo* ^30^. In contrast, neurogenesis is not prominent in the RCs, as TUBIII-immunoreactive neurons were absent in samples initiated at 1 DPI (Figure 3G), and only a few cells resembling immature neuroblasts were sparsely present in those derived from 3 DPI (Figure 3G and Figure S3E), raising the possibility that these may originate from dedifferentiated astrocytes ^20^ or perhaps some few SEZ-derived progenitors introduced in the cultures at this later time point ^22^.

In summary, despite slight variations in oligodendrocyte and neuronal populations, cultures established from both 1 and 3 DPI cortices exhibited similar glial compositions, expansion dynamics, and differentiation patterns. This consistency suggests that initiating cultures within this early post-injury timeframe does not significantly alter the fundamental dynamics of the RC environment. Moreover, our data support that the RCs faithfully recapitulate *in vitro* the *in vivo* injured brain gliotic landscape.

### Proteomic and metabolomic analysis of the RC medium uncovers mediators of brain injury response

Our approach allows direct analysis of the extracellular milieu, including soluble factors and components such as vesicles derived from cell somas, which are difficult to study *in vivo* due to restricted access to this compartment. Thus, we next determined whether the conditioned medium of RCs is representative of gliosis. To address this, we performed proteomic and metabolomic analyses of media from 4 days-cultured RCs established at 1 DPI, and compared it to media from control cells cultured under identical conditions and duration (Figure 4A). The control cells for both proteomic and metabolomic analyses were mouse cortical astrocytes isolated at postnatal day 5 (P5As), known for supporting neuronal survival and health ^31^. For proteomics, we also included human fibroblasts (HFs) as a non-neuronal, non-rodent reference lacking inflammation or neural-specific protein signaling pathways.

**Figure 4.**
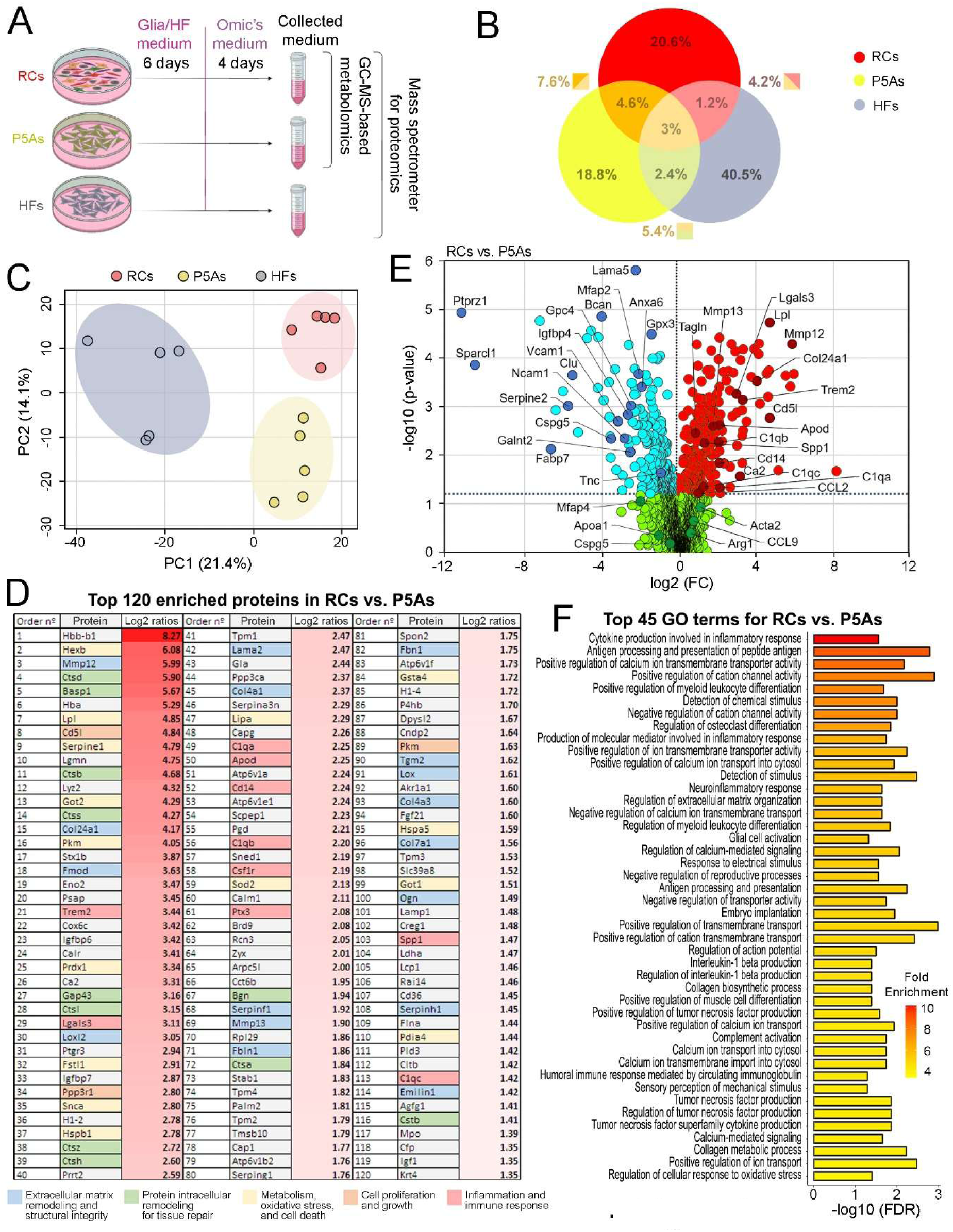
The extracellular medium from RCs resembles a gliosis-like environment. (**A**) Schematic representation of the experimental timeline and omic analysis approaches. Medium from RCs, P5As, and HFs was collected after 6 days of culture in glia or HF medium and 4 additional days in proteomic medium. Metabolites and proteins in the collected medium were analyzed using gas chromatography-mass spectrometry (GC-MS) metabolomics and mass spectrometry-based proteomics. (**B**) Venn diagram illustrating the shared and unique proteins identified in the media of RCs, P5As, and HFs, represented as percentages of the total proteins. (**C**) PCA plot of proteomic profiles shows clear separation among RCs (red), P5As (yellow), and HFs (gray), reflecting distinct proteomic milieus. (**E**) Volcano plot comparing RCs to P5As highlights the differential expression of proteomic markers. The x-axis represents the difference in normalized log2 abundances (RCs - P5As), calculated as the difference in average Zq values between the two conditions. Red circles indicate proteins significantly overrepresented in RCs, while blue circles highlight those with greater representation in P5As (p-value ≤ 0.05). Proteins with non-significant changes are depicted in light green, while those of particular biological interest are highlighted in a darker shade. (**D**) Table presenting the top 120 upregulated genes in RC relative to P5As, represented by a red gradient (standardized log2 ratios integrated by category; p ≤ 0.05, p-value). Genes related to injury response are grouped and color-coded by function, as indicated in the legend. (**E**) Table listing the top 30 GO terms for biological processes enriched in RCs relative to P5As.

Comparison of protein profiles across the three cell culture types revealed that the conditioned media derived from RCs, P5As and HFs were very different (Figure 4B). Of note, a closer similarity between RCs and P5As, sharing 7.6% of total proteins, could be observed. In contrast, RCs and HFs shared only 4.2% and P5As and HFs shared 5.4% (Figure 4B). This pattern aligns with the distinct germ layer origins and species differences among the three groups. Moreover, the pronounced differences between the glial cultures, RCs and P5As, coupled with the higher proportion of proteins shared between HFs and P5As than between HFs and RCs, highlight distinctive characteristics of the RC-derived medium. Principal component analysis (PCA) reinforced these observations, revealing clear separation in protein composition across the three media types, with slightly greater divergence between RCs and HFs than between P5As and HFs (Figure 4C).

To further explore the distinct profile of RCs, we first focused on proteins enriched in the RC-derived medium compared to that from P5As. We identified a set of 411 proteins with significantly different levels (p ≤ 0.05, p-value), of which 230 were increased and 181 decreased. The top 120 enriched proteins list (Figure 4D) and volcano plot analysis (Figure 4E) showed a marked presence of factors involved in extracellular matrix remodeling, such as metalloproteases (MMPs) and collagen proteases (COL24A1), which are essential for matrix reconstruction in the injured brain ^32,33^, and a wide spectrum of inflammation-related proteins, including the immune recruitment chemokine CCL2 and the myeloid receptors CD14 and TREM2, all crucial for immune activation ^34^. Additionally, we identified proteins linked to oxidative stress and energy management, such as Lipoprotein Lipase (LPL) and Apolipoprotein D (APOD) ^35,36^. Intriguingly, GALECTIN-3 (Lgals3), an extracellular glycoprotein mediating dedifferentiation of astrocytes into stem-like cells forming neurospheres following invasive injuries in both mouse ^19^and human contexts ^21^, was also notably enriched in the RC medium (Figure 4D, position 29; Figure 4E, volcano plot). Proteins underrepresented in RCs compared to P5As include components of the normal extracellular matrix in the brain, such as Mfaps, Serpine2, Cspg5, and Tnc, as well as anti-inflammatory molecules, including Gpx3, Clu, and Apoa1 (Figure 4E).

In a subsequent analysis, we compared RC-versus-HFs media (Figure S4A-C) and found increased metalloproteinases, lysosomal proteases, oxidative stress response factors, and immune modulators, many of which overlapped with proteins identified in the RC-versus-P5A comparison (Figure 4; Figure S4A, B). Representative examples include numerous astroglia-derived proteins such as serine peptidase inhibitors linked to epileptic lesions ^37^, apolipoproteins associated with Alzheimer’s disease (AD) ^38^, Phosphoprotein Enriched in Astrocytes 15 (PEA15), widely present in reactive astrocytes ^39^, and Secreted Phosphoprotein 1 (SPP1, also known as osteopontin), which is associated with neurocognitive disorders and neuroinflammation ^40^. Relevant examples of proteins correlated with activated myeloid cells are the LPS receptor CD14, involved in microglial phagocytosis ^41^, and complement components such as C1q. We also found a large spectrum of proteins more specific to macrophages than microglia, such as MMP-12 ^42^ previously mentioned, the chemokine CCL9, which recruits immune cells to injured brain tissue ^43^, and Arginase-1 (ARG1), a hallmark of infiltrating macrophages in lesion environments ^44^. Finally, α-SMA (Acta2) confirmed the presence of pericytes ^29^, while the myelination-associated protein Carbonic Anhydrase II ^45^ supports ongoing oligodendrocyte development in the RCs.

Gene ontology (GO) analyses reinforced the findings from individual protein assessments, highlighting features characteristic of an injured gliotic environment (Figure 4F and Figure S4C). Among the top-ranked categories in the RC versus P5As comparison (Figure 4E), prominent terms are *Cytokine production involved in inflammatory response*, *Neuroinflammatory response*, *Regulation of extracellular matrix organization*, *Glial cell activation*, *Antigen processing and presentation*, *Complement activation* and *Regulation of cellular response to oxidative stress*. Similarly, the RC versus HFs comparison (Figure S4C) revealed enriched categories such as *Cytokine production involved in inflammatory response*, *Antigen processing and presentation of peptide antigen*, *Myelination*, and *Cellular oxidant detoxification*.

Finally, we sought to determine whether the RC medium provides insights into the extracellular metabolic milieu of the injured brain, a task also not feasible *in vivo*. To this end, we compared mediums from the RCs with those from P5As (Figure 4A and Figure S5), identifying individual metabolites with significant differences through univariate analysis and mapping metabolic pathways by quantitative enrichment analysis (QEA). PCA plot initially revealed a clear metabolic separation between the P5As and RCs populations (Figure S5A), highlighting their distinct profiles. A volcano plot (Figure S5B) and heatmap (Figure S5C) highlighted these differences, identifying oxidative stress-related compounds in the RC medium, such as pyroglutamic acid (PGA) ^46^ and urea ^47^. Furthermore, among the top 25 statistically significant metabolic pathways with increased representation, a substantial portion were associated with oxidative stress management and injury-related contexts (Figure S5D). For instance, glutathione synthesis, essential for managing ROS, depends on glycine-serine-threonine and cysteine-methionine metabolism ^48,49^, which are highly represented in RCs. Similarly, lipoic acid and glyoxylate-dicarboxylate pathways indicate heightened antioxidant responses to ROS ^68,69^. Arginine biosynthesis and arginine-proline metabolism are linked to increased nitric oxide production under neuroinflammatory conditions ^50^, and elevated glycolysis and the citrate cycle suggest metabolic shifts and mitochondrial adaptations in response to injury ^51,52^. Likewise, purine metabolism may indicate a broader metabolic reprogramming triggered by pathological states ^53^.

Thus, the overall proteomic and metabolomic data support that RC-derived media align with a post-injury gliotic brain milieu, particularly regarding matrix remodeling, immune response, oxidative stress management, and metabolic changes, clearly differentiating it from media derived from P5As and HFs.

### Reactive glial cultures reproduce cellular responses to injury

Our previous findings indicate that the RC environment closely resembles a gliotic environment in terms of cellular composition and secretome. Furthermore, the capacity of RCs to support oligodendrogenesis suggests they might induce complex biological responses to injury as well. To investigate this possibility, we first evaluated whether RC-derived conditioned medium exerts neurotoxic effects on neurons, as neuronal loss is an early-stage hallmark of acute brain injury ^5^ (Figure 5A). To this aim, we separately cultured reactive glial cells and P5As, allowing secreted factors to accumulate in the medium for 7 days. Next, conditioned media from RCs and P5As culture were transferred to 10-day cultured E14 cortical neurons, and assessed neuron survival 5 hours later (Figure 5A-C). Neurons exposed to RC-conditioned medium showed significant death, evidenced by increased caspase-3 activation and fewer TUBIII⁺ cells, unlike those exposed to P5A-conditioned medium, which showed no obvious signs of decreased viability (Figure 5B, C). These data thus demonstrate that molecules either present in or absent from RC-conditioned medium contribute to a neurotoxic environment, resembling *in vivo* injury conditions.

**Figure 5.**
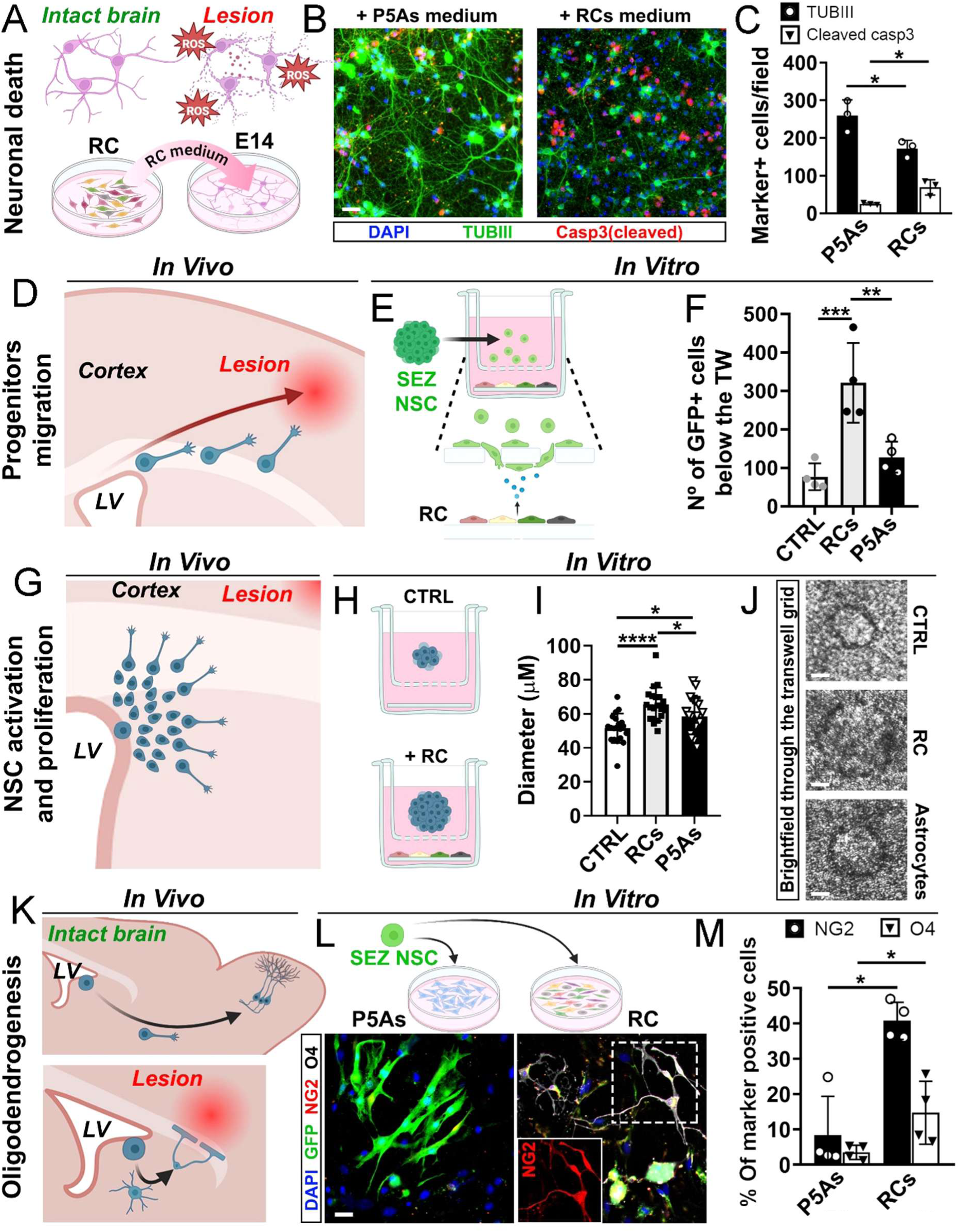
Reactive glial cultures elicit cellular responses to brain injury. **(A)** Schematic of neuronal death following brain injury (top) and the corresponding *in vitro* assay (bottom), where RC-conditioned medium was applied to 10-day-old primary cortical neurons. **(B)** Micrographs of neurons exposed to P5As-(left) or RC-conditioned medium (right) for 5 hours, stained with DAPI, TUBIII, and cleaved caspase-3. Note fewer TUBIII⁺ neurons and increased cleaved caspase-3 in the RCs’ medium condition. (**C**) Histogram confirming the toxic effect of RC-conditioned medium on P5As. (**D**) Schematic of SEZ-derived NSC migration toward the injured cortex *in vivo*. (**E**) Setup to assess NSC migration *in vitro* toward reactive cells, using GFP-labeled ACTb-EGFP SEZ-derived cells plated on 8-μm transwells above RCs. (**F**) Histogram showing significantly higher NSC migration in the RC condition vs. CTRL and P5As. **(G)** Schematic of NSC proliferation in response to cortical injury *in vivo*. **(H)** *In vitro* assessment of this effect, with SEZ-derived NSCs cultured on 0.3-μm transwells above CTRL, RC, or P5As cultures, with neurosphere size after 7 DIV as a proliferation indicator. **(I)** Histogram showing greater neurosphere diameters in RC co-cultures, indicating increased NSC proliferation. **(J)** Brightfield images through the transwell grid of neurospheres showing size differences across conditions. **(K)** Schematic of progenitor cell differentiation into oligodendrocytes at lesion sites vs. olfactory bulb interneuron formation in intact brain regions. **(L)** *In vitro* assay replicating this, with GFP-labeled SEZ-derived neurospheres cultured on P5As (left) or RCs (right) for 7 days in differentiation conditions, where RCs drive NSC differentiation into NG2⁺ (red) and O4⁺ (white) oligodendroglial cells. **(M)** Histogram showing a significantly higher proportion of GFP⁺ oligodendrocytes in RCs compared to P5As. Data in panels C, F, I, and M are mean ± SD (n = 3). Statistical analysis: Student’s t-tests for C and M, and one-way ANOVA with FDR correction for F and I, *p < 0.05; **p < 0.01; ***p < 0.001; ****p < 0.0001. Scale bars: 50 μm.

Aiming to validate the model’s suitability for studying the brain’s regenerative potential, we next examined the behavior of aNSCs or progenitors upon exposure to RCs. To this end, we first assessed whether RCs promote the recruitment of neural progenitors, a response commonly observed following brain injuries *in vivo* ^54^ (Figure 5D). To this aim, we cultured the reactive cells, P5 astrocytes (P5As), or maintained cell-free medium for 7 days. Following this, we added aNSCs derived from cultured SEZ neurospheres onto the culture chamber, separated from the underlying cells by an 8-micron porous transwell, which allows cell migration through the membrane. In these experiments, NSCs from transgenic GFP-Actin mice enabled the identification of their progeny through GFP expression (Figure 5E). After 24 hours, an increased number of GFP⁺ cells (threefold) had migrated in the RC condition compared to the P5As condition or cell-free medium (Figure 5F), demonstrating that RCs release chemotactic signals that attract SEZ-derived neural progenitors, thus echoing the response observed in lesion brain tissues *in vivo*.

We next aimed to explore whether the RC replicates injury-induced aNSC proliferation observed *in vivo* ^55^ (Figure 5G). To do this, we cultured reactive glia, P5As, or maintained cell-free medium as before. After 5 days, we introduced SEZ-derived aNSCs into an upper transwell chamber with a 0.4-micron porous membrane, allowing soluble factor exchange but preventing cell migration, and maintained the co-cultures under proliferative (+EGF/+FGF), non-adherent conditions for 7 days to promote neurosphere formation (Figure 5H). We observed that neurospheres in co-culture with reactive glial cells were significantly larger than with P5As (1.13 fold increase) or cell-free medium (1.27 fold increase), indicating that the RC environment promotes NSC expansion (Figure 5I, J).

Following these observations, we next explored whether RCs influence cell fate decisions and plasticity. First, we examined whether RCs promote selective differentiation of SEZ-derived aNSCs into oligodendrocytes, a phenomenon commonly observed in injured brains ^8,30^ (Figure 5K). To achieve this, we cultured reactive glia or postnatal astrocytes for 7 days. Next, NSCs originating from dissociated neurospheres derived from SEZ of actin-GFP mice were seeded onto pre-established cultures (RCs or P5As) and maintained under differentiation conditions for an additional week (Figure 5L). Following that, we assessed whether GFP⁺ cells differentiated into OPCs or oligodendrocytes by evaluating NG2 and O4 expression. In these experiments, there was a significantly higher proportion of cells expressing either of these markers (4.63-fold increase for NG2; 4.21-fold increase for O4) in the RC condition compared to the P5As condition (Figure 5L, M), thus demonstrating that RCs enhance oligodendrocyte production from aNSCs.

Second, we investigated whether exposing resting adult cortical glia to RCs induces them to form neurospheres *in vitro*, mimicking the behavior observed in astrocytes isolated from the mouse or human brain following injury *in vivo* ^19,21^. To explore this possibility, we dissected cortices from non-injured mice and cultured the dissociated cells either alone or in co-culture with RCs (Figure S6A). In this paradigm, it is important to note that, due to their injury origin, RC-derived cells were potentially capable of forming neurospheres. To account for this, we also cultured reactive cells alone as control. After 7 days under proliferative conditions, co-cultures of uninjured cortical cells with reactive glia generated approximately 15 times more neurospheres than uninjured cells cultured alone and twice as many as RC alone (Figure S6B, C).

Taken together, these findings, provide compelling evidence that our RC model faithfully reproduces a wide range of *in vivo* cellular responses to brain injury.

### Reactive cultures are a powerful tool to explore reactive glia-to-neuron conversion

Neuronal reprogramming *in vivo* has potential for regenerative medicine applications ^56^. However, understanding the conversion process in the injured brain is challenging due to difficulties in targeting specific glial populations with reprogramming vectors and reliably tracking the reprogrammed cells over time. Therefore, we explored whether our culture system could help to address these questions as well. First, we investigated the effects of retroviral combinations encoding the reprogramming factors *Neurog2* and *Bcl2*, *Ascl1* and *Sox2*, or *Neurog2* alone, all of which have demonstrated varying capabilities to reprogram glial cells into iNs *in vivo* ^12,57^. To achieve this, we exposed 3 DPI-derived RCs precultured during 5 days to retroviral vectors encoding the above mentioned genes together with *myc*-epitope and RFP reporters, and assessed neuronal reprogramming 14 days later, based on TUBIII expression in the reporter-positive cells (Figure 6A-D).

**Figure 6.**
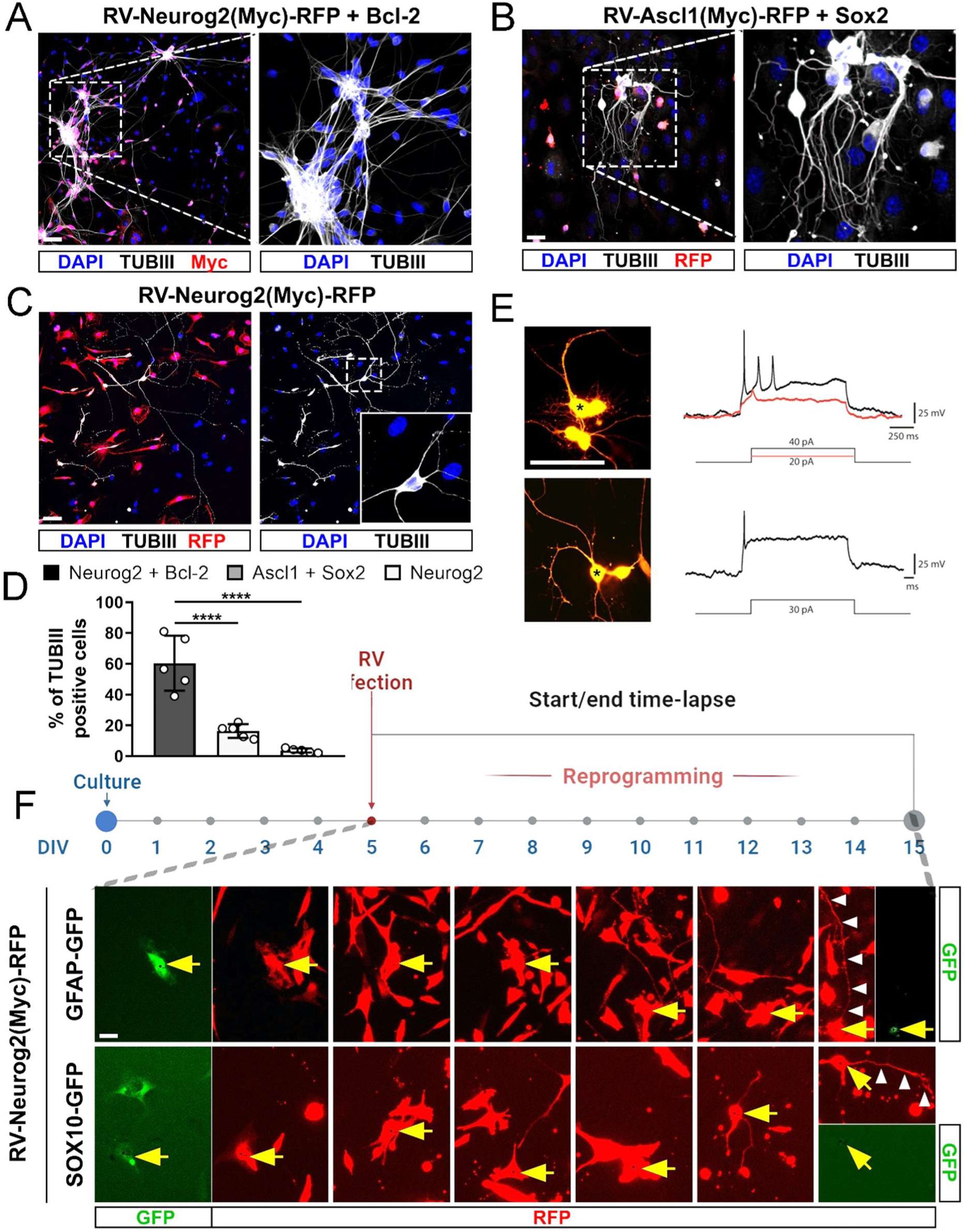
Validation of reactive cultures for neuronal reprogramming and for tracing the reactive glia-to-neuron conversion. (**A–C**) Micrographs showing immunocytochemistry of reactive cultures 14 days post-infection with retroviruses encoding Neurog2 + Bcl-2, Ascl1 + Sox2, or Neurog2 alone, carrying reporters (c-Myc or RFP) as indicated. Among transduced cells identified by reporter expression (red), Neurog2 + Bcl-2 (A) produces the highest proportion of TUBIII⁺ (white) iNs, followed by Ascl1 + Sox2 (B) and Neurog2 alone (C). (**D**) Quantification of TUBIII⁺ cells confirms the reprogramming efficiency hierarchy described in (C). (**E**) Patch-clamp recordings of Neurog2-reprogrammed cells show action potential generation in induced neurons (iNs). (**F**) Time-lapse images of reactive cultures transduced with RV-Neurog2-RFP illustrate glial-to-neuron conversion over time, accordingly with the timeline above. Cultures from GFAP::GFP and Sox10::GFP transgenic mice identify astroglial (GFP⁺) and oligodendroglial (GFP⁺) origins, respectively. Yellow arrows indicate cells transitioning to neuronal morphology, with development of elongated processes (white arrowheads) and progressive GFP signal loss from glial markers (GFAP or Sox10). Scale bars: 100 µm (A–E), 30 µm (F). Data in (D): mean ± SEM; statistical analysis One-way ANOVA, ****p < 0.0001.

The results revealed a hierarchy of reprogramming efficiencies mirroring previous *in vivo* findings, with *Neurog2* and *Bcl2* yielding the highest conversion rate up to 64% (Figure 6A, D) compared to ∼75% *in vivo* ^57^. *Ascl1* and *Sox2* resulted in 21% conversion rate (Figure 6B, D) compared to ∼30% *in vivo* ^12^, while Neurog2 alone had a modest 4% efficiency (Figure 6C, D), slightly higher than ∼2% *in vivo* ^57^.Patch-clamp analysis confirmed that iNs could generate action potentials, even those generated upon forced expression of *Neurog2* alone (Figure 6E). Therefore, our data confirm that RC-derived cells can be reprogrammed into neurons with efficiencies comparable to *in vivo* conditions, validating the model for studying reprogramming mechanisms.

Building on these results, we next explored whether the RC model could help to identify adult glial lineages amenable to neuronal conversion. To this aim, we performed genetic fate-mapping studies. For these experiments, RCs were derived from injured tissues isolated at 3 days following stab wound injury performed in hGFAP::GFP mice and Sox10::GFP mice. We tracked GFP^+^/RFP^+^ cells from RC transduced with a retrovirus encoding *Neurog2(RFP)* by monitoring GFP and RFP fluorescence over 10 days using time-lapse video imaging (Figure 6F). This allowed us to identify whether iNs derived from reactive astrocytes or cells belonging to the oligodendroglial lineage. Still images from Figure 6F and corresponding to the Movies S1 and S2 show GFP-labeled astrocytes and oligodendrocytes transitioning into neuron-like shapes with characteristic processes. Notably, GFP levels decreased over time, while the cells ceased expressing GFAP or Sox10 as they transitioned into neurons (Figure 6F, Movies S1 and S2).

Collectively, these findings establish the RC as a powerful tool for monitoring the reprogramming of glial cells into neurons. Looking ahead, future studies could refine this approach by integrating time-lapse methodologies with proliferation and morphological analyses, or by using endpoint immunostaining with neuronal markers, as we did in other reprogramming paradigms ^57^.

## DISCUSSION

In this study, we present a novel culture model specifically developed to investigate the mechanisms underlying brain injury and its impact on aNSCs behavior and glial lineage plasticity. Initially developed to investigate the acquisition of stem cell potential by adult reactive astrocytes ^19,22^, the model primarily focused on astroglial populations. Here, we broaden the scope of the model, demonstrating that RCs exhibit a striking cellular diversity and dynamic properties that closely emulate the gliotic milieu of the injured brain, including fostering glial and NSCs behaviors characteristic of its reactive environment in vivo. Thus, RCs provide a robust platform to study mechanisms underlying brain injury. Furthermore, our approach enables sampling of the soluble extracellular compartment for proteomic and metabolomic analyses, which is not feasible *in vivo*.

Two key factors were essential for the development of the RCs. First, glial cells were activated *in vivo* using a refined and specialized stab wound-based method, which we identify as critical to the success of the RCs. According to our data, this method effectively replicates the temporal progression of gliosis described by Frik et al. ^58^, with activated myeloid cells appearing first, followed by reactive macroglia (Figure 1D). At early stages of establishment, the cellular composition of RCs closely resembles that observed *in vivo* at 3 DPI (Figure 2D), supporting the notion that the cultured cells undergo a natural activation process. Further evidence for this is provided by the hallmark features of *in vivo* injuries exhibited by RCs, including CD68 and CD11b expression in myeloid cells ^28^ and GFAP in astrocytes, as well as a robust profile of proteins and metabolites associated with gliosis in their secretome. Consequently, our method prevented the need for artificial compounds to induce gliosis as required by others ^17,59^. Importantly, our findings indicate that cultures established at 1 or 3 DPI can be used almost interchangeably, as they show no major differences in cell composition, except for the further progression of oligodendrogenesis (Figure 3H) and the presence of SEZ-derived progenitors in the later ^22^.

The second key factor is an intrinsic effect of glial reactivity, which enhances cell adhesion, allowing rapid substrate attachment and facilitating a seamless transition from *in vivo* to *in vitro* conditions within a minimal time frame. Notably, this increased adhesion affects all cell types, and thus, RCs result in a multilineage milieu that includes the most representative cellular lineages of the injured brain, in contrast to FACS-or immunomagnetic-based methods, which typically target specific ones^60–62^. Although cellular adhesive changes following injury have been studied for decades ^63^, the molecular mechanisms underlying the rapid attachment observed in our approach remain unclear and could be further investigated using the RC model itself. Supporting this, our proteomic analysis identified numerous cell adhesion-related proteins in the RC secretome, such as MMPs ^32^ and osteopontin ^64^, among others.

Apart from adhesion enhancement, RCs recapitulate a broad spectrum of injury-related processes, thereby allowing the investigation of other multiple aspects of brain pathology. In this sense, integrating the cellular responses to injury reproduced by RCs with proteomic and metabolomic data could significantly aid in identifying underlying molecular mechanisms. For example, neuronal death induced by the RC may be driven by cathepsins (Cts; Figure 4D), which are significantly enriched in its conditioned medium and are known to play a role in neuronal toxicity observed in diseased brains *in vivo* ^65^. In addition, we identified oxidative stress-associated metabolites, such as pyroglutamic acid (PGA) ^46^ and urea ^47^, in the RC medium. These findings align with the activation of an antioxidant response, as indicated by enhanced cysteine metabolism detected through metabolomic analysis. ^48^. Notably, the RC medium contains other proteins highly relevant to neurodegenerative diseases, such as alpha-synuclein (Snca) ^66^.

Regarding cellular behavior effects, we have specifically focused on how RCs influence stem cell behavior and the acquisition of multipotency by glial cells, given their potential role in regenerative processes. In this context, omics data provide insights into the clonal expansion of SEZ-derived NSCs following injury *in vivo* ^55^ and their migration toward the lesion site ^54^, both processes faithfully recapitulated by RCs (Figure 5). For instance, IGF-1, IGF-2, and Progranulin (GRN) may be responsible for promoting proliferation of neural progenitors ^67,68^, while acyl-CoA-binding protein (ACBP or DBI) may drive to NSC expansion ^69^, when co-cultured with RCs. Furthermore, cell proliferation is consistent with the enhancement of glycolytic metabolism ^70^ identified through metabolic analysis of the cell secretome (Figure S5). Regarding migration, the RC medium showed enrichment of CXCL12, a chemokine known to recruit NSCs to injury sites following stroke ^71^. But this process can also involve SLIT2, which promotes neural progenitor guidance and proliferation via ROBO receptors ^72^, and neuropilins, which mediate semaphorin-dependent chemotaxis ^73^.

Among all the injury responses reproduced by RCs, perhaps one of the most striking is their influence on fate decisions and plasticity, specifically promoting the differentiation of neural progenitors toward an oligodendroglial lineage and compelling mature glial cells from non-neurogenic niches to form neurospheres. Our proteomic analysis has also identified candidates here, such as GALECTIN-3, which promotes oligodendrogenesis ^74^ and dedifferentiation of reactive astrocytes into stem cell-like cells ^19,21^. Other relevant candidates in this context are IGF1 and its receptor IGF1R, which are involved in OPCs proliferation, differentiation toward oligodendrocytes ^75^ and maturation ^76^.

A last principal feature of RCs is that they derive from adult tissue, which aligns better with pathophysiological contexts than models using postnatal or embryonic tissue sources ^59^. This also provides a strong foundation for exploring the effects of certain manipulations difficult to achieve in the *in vivo* adult brain, including neuronal reprogramming for regenerative purposes. In this field, technical hurdles such as non-specific viral tropism, promoter leakage, and vector silencing pose significant barriers to precisely identifying the glial cells targeted for reprogramming and hinder their tracking throughout the conversion process. Such challenges may have led to misinterpretation of results, sparking debates about whether iNs are consistently genuine reprogrammed cells or preexisting neurons that have been mistakenly identified ^77^. Our data support the potential of RCs to effectively address these issues as well, as they demonstrated reprogramming efficiencies equal to those observed *in vivo* and enabled the tracking of glial-to-neuron conversion through time-lapse videos (Figure 6, see also ^57^). Looking ahead, further studies could investigate whether iNs persist long-term and mature or undergo cell death as observed *in vivo*, and determine whether ferroptosis inhibitors like vitamin E can enhance reprogramming efficiency and iN viability within the RC environment as observed *in vivo* ^57^.

In conclusion, our method offers a simple, robust, and versatile approach to investigate a wide range of cellular processes within the brain injury context. Future research could build upon our proteomic and metabolomic analyses to uncover additional insights, or explore other components in the RC secreted medium, such as transfer RNAs or secretory vesicles. Moreover, applying similar approaches to other models of brain disorders, like Alzheimer’s disease and stroke, could extend the model’s applications to additional pathological frameworks.

## MATERIALS AND METHODS

### Animals

All experiments were performed on adult male mice (8–12 weeks old), primarily C57BL/6J specimens acquired from the Animal Facility at the Universidad Autónoma de Madrid (Registration Number ES28079000018). For the time-lapse video reprogramming experiments, we used transgenic lines, including strains in which enhanced green fluorescent protein (eGFP) is driven either by the human GFAP promoter [TgN(hGFAP-EGFP)GFEA ^78^], or the Sox10 promoter [Tg(Sox10-EGFP)GE255Gsat; Jackson Laboratories, catalog. No. 011876-UCD]. To identify infiltrating immune cells in RCs, adult knock-in mice expressing red fluorescent protein (RFP) under the Ccr2 promoter (Jackson Laboratories, catalog No. 017586) were used in some experiments. For NSC migration and differentiation into oligodendrocytes, transgenic B6 ACTb-EGFP mice (Jackson Laboratories, catalog No. 003291) were used.

All experimental procedures complied with Spanish (license Proex 1.8/21) regulations, as well as European Union directives (86/609/EEC, 2003/65/EC, 2010/63/EU, RD1201/2005, RD53/2013). Measures were rigorously implemented to minimize animal suffering and reduce the number of animals used in each procedure.

### Stab Wound Injury

Stab wound lesions were induced in the cerebral somatosensorial-visual cortex of adult male mice using a modified version of the method we previously reported ^12,79^. Briefly, animals were anesthetized intraperitoneally with fentanyl (0.05 mg/kg), midazolam (5 mg/kg), medetomidine (0.5 mg/kg), and saline, and subsequently positioned in a stereotactic frame. Surgical lancet injuries were performed based on Bregma coordinates: anteroposterior −1.6 mm to −2.5 mm, mediolateral ±1.5 mm to ±2.5 mm, and dorsoventral 0.5 mm. The lancet was moved approximately 1 mm back and forth along the anteroposterior axis to create six incisions, followed by six additional along the mediolateral axis (see Figure 1A and Supplementary Video 1). Along the anteroposterior axis, the blade moves parallel to its edge, resulting in precise cuts. Conversely, mediolateral movement orients the blade perpendicularly to its edge, producing shallow, scratch-like lesions rather than deep cuts (Supplementary Video 1). This standardized approach enables reproducible lesion placement in either unilateral or bilateral hemispheres (see Figure 1A). Anesthesia was reversed using a solution of buprenorphine (0.102 mg/kg), atipamezole (2.5 mg/kg), and flumazenil (0.5 mg/kg). Mice were sacrificed at 1, 3, 5, or 10 DPI.

### Reactive Glia Cell Culture

After dissecting the tissue as described in the *Results* section, the injured cortex was minced into small fragments and collected in HBSS dissection medium (ThermoFisher, 24020117) supplemented with 10 mM HEPES (ThermoFisher, 15630-049). Tissue fragments were digested in a pre-warmed enzyme solution containing 124 mM NaCl, 5 mM KCl, 3.2 mM MgCl₂, 0.1 mM CaCl₂, 26 mM NaHCO₃, 10 mM D-glucose, 0.2 mg/mL kynurenic acid, 0.67 mg/mL hyaluronidase type IV (Sigma, H3884), and 1.33 mg/mL trypsin (ThermoFisher Scientific). This mixture was incubated at 37°C for 15 minutes, followed by mechanical dissociation through gentle pipetting.

Dissociated cells were then plated onto 12 mm poly-D-lysine (PLL)-coated glass coverslips in 24-well plates at one hemisphere per well. Cells were cultured in DMEM/F12 medium supplemented with glucose (45%, 1:100), penicillin/streptomycin (1:100), B27 (1:200), 10% fetal bovine serum (FBS), and GlutaMax. Cultures were maintained for 6 days at 37°C in an 8% CO₂ atmosphere under normoxic conditions. Notably, cells from the lesioned cortex adhered robustly within the first 2 hours, in contrast to cells from non-lesioned or ipsilateral cortices, which displayed minimal or no adhesion (Figure 2A-Cell culture and Figure 2B).

### Mouse Cortical Astrocyte Culture

Mouse cortical astrocytes were cultured as previously described ^12^. Briefly, five-day-old postnatal mice were sacrificed by decapitation. Cerebral cortices were separated under a stereoscope, meninges were removed, and the tissue was dissociated in ice-cold HBSS. Cells were centrifuged at 1400 rpm for 5 minutes at 4°C and transferred to a T-75 flask with astrocyte medium consisting of DMEM F12, 10% FBS, B27 (1:200), Penicillin/Streptomycin (1:100), EGF, and FGF2 (both at 50 ng/ml). Cells were cultured at 37°C, 8% CO₂.

### E14 Cortical Neuron Cultures and experiments

Embryonic cortical neurons were cultured as described by Heinrich and collaborators ^80^. For neurotoxicity assays, cortical neurons were cultured for 10 days before completely removing neuronal media. One milliliter of pre-heated media from 7DIV reactive cultures and astrocytes (stored at −80°C) was added into each well. Neuronal cultures were fixed with 4% PFA for 10 minutes, 5 hours after medium replacement.

### SEZ Neurosphere Cultures

Wild-type and ACTb-EGFP male mice were sacrificed with pentobarbital, and the SEZ-SVZ lining the lateral ventricles was dissected under a stereoscopic microscope (coronal section at the level of the optic chiasm). Tissue was placed in a 6-cm dish containing DMEM-F12, cut into 1 mm pieces, and transferred to a dissociation solution [124 mM NaCl, 5 mM KCl, 3.2 mM MgCl₂, 0.1 mM CaCl₂, 26 mM NaHCO₃, 10 mM D-glucose, 0.2 mg/mL kynurenic acid, 0.67 mg/mL hyaluronidase type IV, 1.33 mg/mL trypsin]. After a 30-minute incubation at 37°C, a trypsin inhibitor solution (1 mg/mL) was added, and cells were mechanically dissociated. The suspension was centrifuged at 1500 rpm for 7 minutes, and the pellet was resuspended in neurosphere culture media (DMEM-F12 + Glutamax, B27 (1:100), Penicillin/Streptomycin 1:100, FGF2 and EGF 20 ng/mL each). Cells were cultured in 24-well plates at a density of 100,000 cells/well, maintained at 37°C in humidified 5% CO₂–95% air incubators for 7 days, replacing one-quarter of the medium every two days.

For progenitor activation and proliferation assays, neurospheres were cultured in 0.3-micron porous transwells on pre-coated PDL coverslips with reactive cultures, P5As, or without cells. Neurospheres were cultured for 7 days at 37°C, 8% CO₂, with one-third of the medium changed every 2-3 days. On day 7, neurospheres were quantified and photographed for diameter measurements using a PAULA (Personal Automated Lab Assistant) microscope. Measurements were performed with ImageJ software.

For migration assays, we use an adaptation of the Boyden chamber assay. Basically, 14-days-old-SVZ-derived neurospheres were isolated by centrifuging them at 1400 rpm for 5 minutes. After disaggregated them enzymatically, neurospheres were incubated with 0,125% trypsin (7 min, 37 °C) and afterwards, trypsin activity was inhibited by adding FBS. After centrifuging at 1400 rpm for 5 minutes, cells were resuspended with differentiation medium (containing: DMEM F12, B27 (1:100) and Penicillin/Streptomicin (1:100)) and plated in 8-micron porous transwell at a density of 25.000 cells per transwell insert. NSCs were allowed to migrate for 48 hours. Afterwards, the attached cells in the top of the transwell were trypsinized and removed, and transwells were transferred to new wells of a 24-well plate, each containing 700 µL of 0.15% trypsin to achieve the detaching of the migrated cells. Solution was blocked with 250uL of Fetal Bovine Serum and, after centrifuging the plate 3 minutes at 1200 rpm, green cells were quantified with PAULA microscope (PAULA Smart Cell Imager, Leica Microsystems).

For NSCs differentiation assays, GFP SEZ-derived neurospheres were obtained as previously described. After 14 DIV, isolated neurospheres were disaggregated both mechanically and enzymatically (0.125% of trypsin for 7 minutes), and isolated cells were cocultured with 7 DIV reactive cultures or astrocytes with differentiated medium (DMEM F12 (½), Neurobasal (½), Glutamax (1:200), B27 (1:100), penicilin/streptomycin (1:100)). NSCs were plated at a density of 85.000 cells/well. Differentiation process was maintained for 7 days and then, cultures were fixed with 4% PFA for 10 minutes.

### Reprogramming Experiments

Retroviral transduction of reactive glial cultures was performed 5 days after initial plating from 3 DPI mouse brain tissue. Retroviruses encoding reprogramming factors Neurog2 + Bcl-2, Ascl1 + Sox2, or Neurog2 alone were used, with each vector carrying either a myc-epitope tag or RFP reporter under the control of an internal chicken β-actin promoter with cytomegalovirus enhancer (pCAG). Viral particles were generated using VSV-G pseudotyping and titered by clonal analysis to ensure consistent infection efficiency.

Transduction was carried out by incubating the cultures with viral supernatants for 2–3 hours, after which the medium was replaced with a fresh differentiation medium consisting of DMEM/F12 supplemented with B27, 3.5 mM glucose, penicillin/streptomycin (1:100), and buffered with HEPES. Cultures were maintained for 14 days post-transduction to assess reprogramming efficiency by immunocytochemistry for TUBIII expression.

For lineage tracing, reactive cultures derived from GFAP::GFP or Sox10::GFP transgenic mice were infected with RV-Neurog2-RFP, allowing for the identification of astroglial and oligodendroglial origins, respectively. Time-lapse imaging was performed for 10 days post-transduction, tracking changes in GFP and RFP fluorescence and the development of neuronal morphology characterized by elongated processes.

### Time-Lapse Video Microscopy

Time-lapse imaging of reactive cultures was conducted using a cell observer microscope system (Zeiss) at a constant temperature of 37°C with 5% CO2. Phase contrast images were captured every 5 minutes, while fluorescence images were acquired every 8 to 12 hours for a total duration of 10 days. Imaging was performed using a 20x phase contrast objective (Zeiss) equipped with an AxioCamHRm camera. For tracking individual cell conversion events, custom software was used for automated tracking and analysis. Video sequences were assembled using ImageJ software (National Institutes of Health, USA) and played back at 30 frames per second to visualize glial-to-neuron conversion over time.

### Electrophysiological Studies

Whole-cell patch-clamp recordings were performed at room temperature to evaluate the electrophysiological properties of reprogrammed cells. Micropipettes were made from borosilicate glass capillaries, with electrodes having resistances of 2–2.5 MΩ. Pipettes were tip-filled and back-filled with internal solution containing 136.5 mM K-gluconate, 17.5 mM KCl, 9 mM NaCl, 1 mM MgCl2, 10 mM HEPES, and 0.2 mM EGTA (pH 7.4, 300 mOsm). The external solution consisted of 150 mM NaCl, 3 mM KCl, 3 mM CaCl2, 2 mM MgCl2, 10 mM HEPES, and 5 mM glucose (pH 7.4, 310 mOsm).

Reprogrammed cells were selected for recordings based on RFP fluorescence. Signals were sampled at 10 kHz and filtered at 5 kHz using an Axopatch 200B amplifier (Axon Instruments). Action potentials were induced by step-current injections, and input resistance was calculated from hyperpolarizing current injections at a holding potential of −70 mV.

### Media collection for proteomic and metabolomic analysis

Reactive cultures were established as previously described. Mouse cortical astrocytes were cultured as outlined above. Simultaneously, human fibroblasts from a 41-year-old male control donor were expanded in T-75 flasks with media containing DMEM, 10% FBS, Penicillin/Streptomycin (1:100), and EGF (1:5000). Once both astrocytes and fibroblasts reached 80% confluence (2-5 days in culture), cells were plated in a 24-well plate with PDL-coated coverslips at a density of 75,000 astrocytes and 55,000 fibroblasts per well. After reaching 80% confluence in 24-well plates, similar to reactive cultures after 7 days, cells were washed twice with pre-warmed proteomic media (DMEM without phenol red, Glutamax 1:100, N2 1:50, Penicillin/Streptomycin 1:100). Fresh proteomic media (1.3 mL) was added per well, and cells were incubated for 96 hours. Culture media was then collected, and total protein content was quantified by Bradford Assay (Bio-Rad). Media were stored at −80°C for subsequent proteome and metabolome assays.

### Proteomic Analysis

To prepare samples for proteome analysis, secretome proteins were denatured in lysis buffer containing 50 mM Tris-HCl (pH 7.8), 2% SDS, and 5 mM TCEP. Samples were boiled for 5 minutes, allowed to cool to room temperature, and incubated for 20 minutes. The denatured proteins were then centrifuged at 15,000 g for 10 minutes, and protein concentrations were measured using the RCDC Protein Kit (BioRad). Protein digestion was carried out using the filter-aided sample preparation (FASP) method (Wiśniewski et al., 2009) with 30 kDa centrifugal filter devices, followed by overnight digestion with modified trypsin (Promega) at a protein-to-trypsin ratio of 50:1 at 37 °C. Peptides were subsequently desalted using Oasis HLB columns (Waters) before labelling.

For isobaric labelling, tryptic peptides were dissolved in 150 mM TEAB buffer, and peptide concentration was measured with the Direct Detect system (Merck Millipore). Equal amounts of peptides from each sample were labelled using TMTpro 18plex reagents (Thermo Fisher), with modifications to the manufacturer’s protocol. Labelling reaction was terminated, after a 1-hour incubation, by adding 0.5% trifluoroacetic acid (TFA) and incubating for 10 minutes. Labeled peptides were combined, and excess reagents were removed by desalting with Oasis HLB columns.

The labeled peptide samples were then suspended in Buffer A (0.1% formic acid), loaded onto Evotip C18 microcolumns (Evosep Biosystems), and separated on an Evosep One chromatography system coupled to a Thermo Orbitrap Eclipse mass spectrometer. Peptide separation was achieved using an 88-minute gradient with 15 samples per day (SPD) pre-programmed gradients. The Orbitrap Eclipse was set to acquire enhanced Fourier transform resolution spectra (120,000 resolution) within an m/z range of 375–1,500, followed by MS/MS spectra of the most intense ions. High-energy collisional dissociation (HCD) fragmentation was applied with a collision energy of 36, targeting 50,000 ions at a resolution of 30,000, and with dynamic exclusion set to 30 seconds.

Mass spectra were processed and searched against the mouse Uniprot database (June 2022) using the Sequest HT algorithm in Proteome Discoverer 2.5, with decoy sequences included to estimate false discovery rates. Search parameters included trypsin as the enzyme, allowing up to two missed cleavages, with mass tolerances of 2 Da for precursors and 0.03 Da for fragments. Fixed modifications included carbamidomethylation of cysteine (+57.021), while variable modifications included TMTpro labelling (+304.207), methionine oxidation (+15.995), and N-terminal acetylation (+42.011) of proteins.

### Metabolomic Analysis by Gas Chromatography–Mass Spectrometry

Metabolites were extracted from 50 μL of medium by adding 250 μL of acetonitrile containing 7 ppm 4-nitrobenzoic acid as an internal standard. The mixture was vortex-mixed briefly and shaken at 100 rpm for 10 minutes at 4 °C to ensure thorough mixing. After incubation, samples were centrifuged at 12,000 rpm for 20 minutes at 4 °C. Supernatants (120 μL) were transferred into GC-MS vials and dried using a SpeedVac concentrator. For derivatization, dried extracts underwent oximation by adding 20 μL of O-methoxyamine hydrochloride in pyridine, vortex-mixing for 5 minutes, and incubating in darkness at room temperature for 16 hours. Subsequently, trimethylsilylation was performed by adding 20 μL of N,O-bis(trimethylsilyl)trifluoroacetamide (BSTFA) with 1% trimethylchlorosilane (TMCS), vortex-mixing for 5 minutes, and incubating at 70 °C with constant shaking at 400 rpm for 1 hour. Samples were then cooled to room temperature for 1 hour. Finally, 100 μL of heptane containing 5 ppm methyl stearate (external standard) was added, and the samples were centrifuged at 2,000 rpm for 10 minutes at 10 °C prior to GC-MS analysis.

Blanks and quality control (QC) samples were prepared following the same extraction and derivatization procedures. QC samples were created by pooling 20 μL aliquots from each study sample, thoroughly mixing, and dividing into nine separate QC samples. These QCs were analyzed at the beginning, at the end, and after every six samples during the GC-MS run to monitor analytical performance and ensure data quality.

For GC-MS analysis, 1 μL of each derivatized sample was injected in split mode (split ratio 5:1) onto a Restek Rxi-5ms column (30 m × 0.25 mm i.d. × 0.25 μm film thickness) using an Agilent 8890 GC system coupled to a LECO BT GC-TOF-MS. Helium was employed as the carrier gas at a constant flow rate of 1 mL/min. The oven temperature program was as follows: initial hold at 60 °C for 1 minute; ramp at 10 °C/min to 300 °C; hold at 300 °C for 4 minutes. The inlet and transfer line temperatures were set at 250 °C and 280 °C, respectively. Mass spectrometry was conducted in electron impact ionization mode at 70 eV, with an ion source temperature of 300 °C. Data were acquired in full scan mode over an m/z range of 50–600 at a rate of 12 scans per second, with a solvent delay of 5.5 minutes. The total chromatographic run time for each sample was 35 minutes.

Data processing was performed using LECO ChromaTOF Sync software (Version 1.1.5.1). Metabolites were identified by matching deconvoluted spectra with the NIST 2020 Mass Spectral Library (version 2.4). The abundance of each metabolite was normalized to the internal standard (4-nitrobenzoic acid), and compounds with higher abundances in blank samples were excluded to eliminate potential contaminants. Statistical analyses, including univariate comparisons, principal component analysis (PCA), volcano plots, heatmaps, and quantitative enrichment analysis (QEA), were conducted using MetaboAnalyst 6.0 to identify significant metabolic differences between sample groups.

### Microscopy and Image Analysis

Microscopy and imaging were performed using a combination of bright-field and fluorescence microscopy. Analysis of the number of neurospheres was conducted using a Nikon Eclipse TS100 bright-field microscope. Fluorescent neurosphere images were obtained using a Leica PAULA (Personal Automated Lab Assistant) microscope. Immunocytochemistry images were acquired with a Nikon Eclipse E600 epifluorescence microscope equipped with filters from Semrock (UV-2A, GFP, YFP, mCherry, Cy5) and a Nikon DS-camera Qi2, along with NIS-Elements software for capturing images. Confocal imaging of brain slices was performed using a Leica TCS-SP5 confocal microscope. All images were subsequently analyzed using Fiji Image J software. The analysis of reactive cell cultures was performed by capturing mosaic images of 64 representative fields using a Leica Thunder Imager Tissue microscope with a 20x objective.

### Data Analysis and Statistical Procedures

Quantifications were conducted manually and assisted by QuPath software (version 0.5.1) for digital image analysis. GraphPad Prism8 Software was used for statistical analysis and graph creation. To compare means between conditions, one-way ANOVA with post-hoc Tukey test or False Discovery Rate (FDR) or Student’s t-test were used. Significant differences are shown as follows: * p<0.05, ** p<0.01, *** p<0.001, **** p<0.0001.

For proteomic data, quantitative analysis was performed using the iSanXoT program. The false FDR was calculated using the corrected Xcorr score, with additional filtering for a precursor mass tolerance of 10 ppm. Identified peptides had an FDR of 1% or lower. Quantitative data were integrated from the spectrum level to the protein level using the WSPP statistical model and the Generic Integration Algorithm (GIA). Differences in peptide and protein abundance were considered significant with p-values < 0.05 using a two-sided t-test.

For metabolomic analysis, statistical analyses included univariate comparisons in Excel to identify significant differences between RC and P5A samples. Multivariate analyses, such as principal component analysis (PCA), volcano plots, heatmaps, and quantitative enrichment analysis (QEA), were performed using MetaboAnalyst 6.0 ^81^.

## ACKNOWLEDGMENTS

We gratefully acknowledge Francisco Sánchez-Madrid and Danay Cibrian (UAM) for kindly providing the transgenic ACTb-EGFP mice. We acknowledge the Omics Core Facilities of the Cajal Institute – CSIC and Centro Nacional de Investigaciones Cardiovasculares (CNIC) for the proteomics and metabolomics analyses. This work was supported by the Spanish Ministry of Science and Innovation (MICINN) (grants RTI2018-099345-B-I00 and PID2021-128796OB-I00; CNS2022-135759) and the Ramón y Cajal Programme (RYC-2015-19185) to SG. DMA was supported by CONAHCYT postdoctoral fellowships (#770727 and #801009). Created in BioRender. Gallego, L. (2025): https://BioRender.com/s07a814; *<u>d01z959</u>*; *<u>r49l216</u>; <u>a06h786</u>*; *<u>https:z34t712</u>*; *<u>o46e268</u>*; *<u>b47o575</u>*; *<u>r44z351</u>*; *<u>k10n446</u>*; *<u>t87b645</u>*; *<u>s98b102</u>*.

## AUTHOR CONTRIBUTIONS

V.A.S., D.M.A., A.V.P.-S., L.G., C.H., B.B., and S.G. performed experiments. E.M. and F.O. carried out specific experimental procedures. J.A.L., A.F., A.M.A., and G.P.F. conducted the omics analysis. V.A.S., D.M.A., A.V.P.-S., L.G., F.O., C.H., S.G., and C.L.-S. analyzed data. C.H. and S.G. conceptualized the model. M.G. and S.G. provided critical resources for the successful execution of the project. R.B. and A.F. provided essential materials. S.G. conceptualized the study and wrote the manuscript. All authors discussed and reviewed the manuscript.

