## Supplementary Figures for "Modeling the acutely injured brain environment *in vitro*"

### Supplementary Figures and Videos.

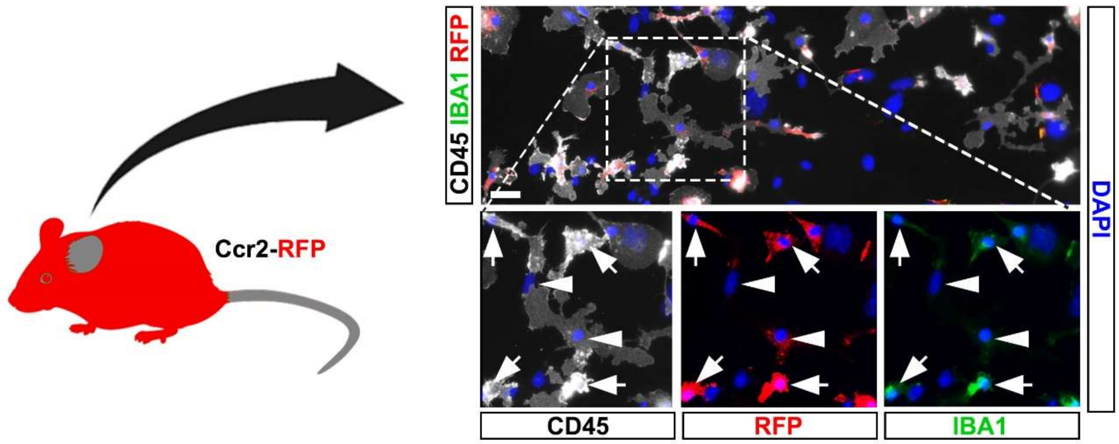


**Figure S1. Ccr2-RFP mice reveal the presence of blood-derived myeloid cells in reactive cultures.** Schematic representation of the Ccr2-RFP mouse model (left), showing red fluorescent protein (RFP) expression under the control of the Ccr2 promoter, which is commonly expressed by blood-derived myeloid cells. Representative fluorescence microscopy images (right) of reactive glial cultures established at 3 days post-injury (DPI) and maintained *in vitro* for 4 days (DIV), showing CD45 (grey), RFP (red), and IBA1 (green) staining, with DAPI (blue) labeling nuclei. Insets display magnified views of the boxed region in the low-magnification image, with white arrows indicating RFP+ cells co-expressing CD45 and IBA1, identifying them as monocyte-derived macrophages within the gliotic environment. Scale bar: 50 μm.

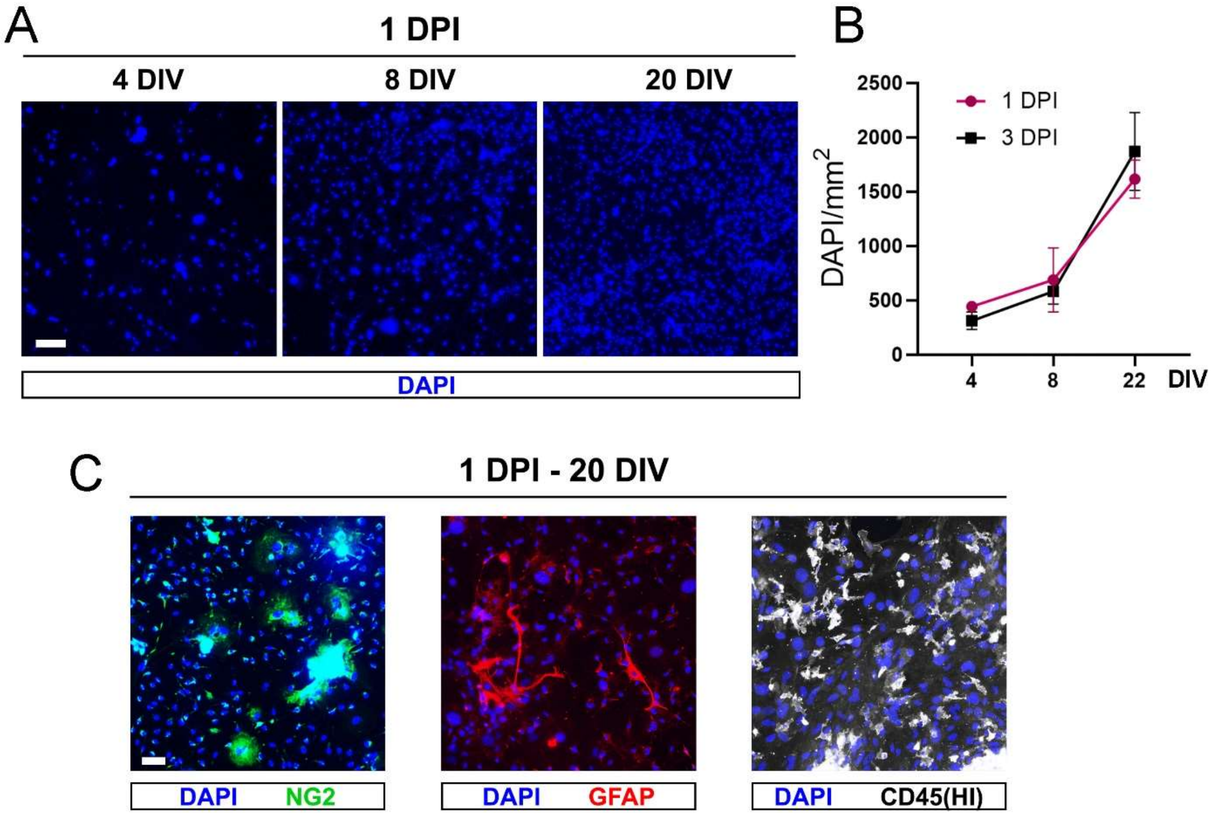


**Figure S2. Temporal dynamics and cellular composition of reactive cultures at different time-points. (A)** Representative fluorescence microscopy images of DAPI staining showing cell density in RCs at 4, 8, and 23 DIV. **(B)** Quantification of DAPI⁺ nuclei per mm² in RCs derived from 1 DPI and 3 DPI lesions, showing time-dependent increase in cell density. **(C)** Representative fluorescence microscopy images illustrating distinct cell populations in RCs at 20 DIV. Staining highlights NG2⁺ oligodendrocyte progenitors (green), GFAP⁺ astrocytes (red), and CD45⁺ (HI: high exposition) immune cells (white). DAPI (blue) marks nuclei in all panels. Error bars represent mean ± SEM. Scale bars: 50 μm.

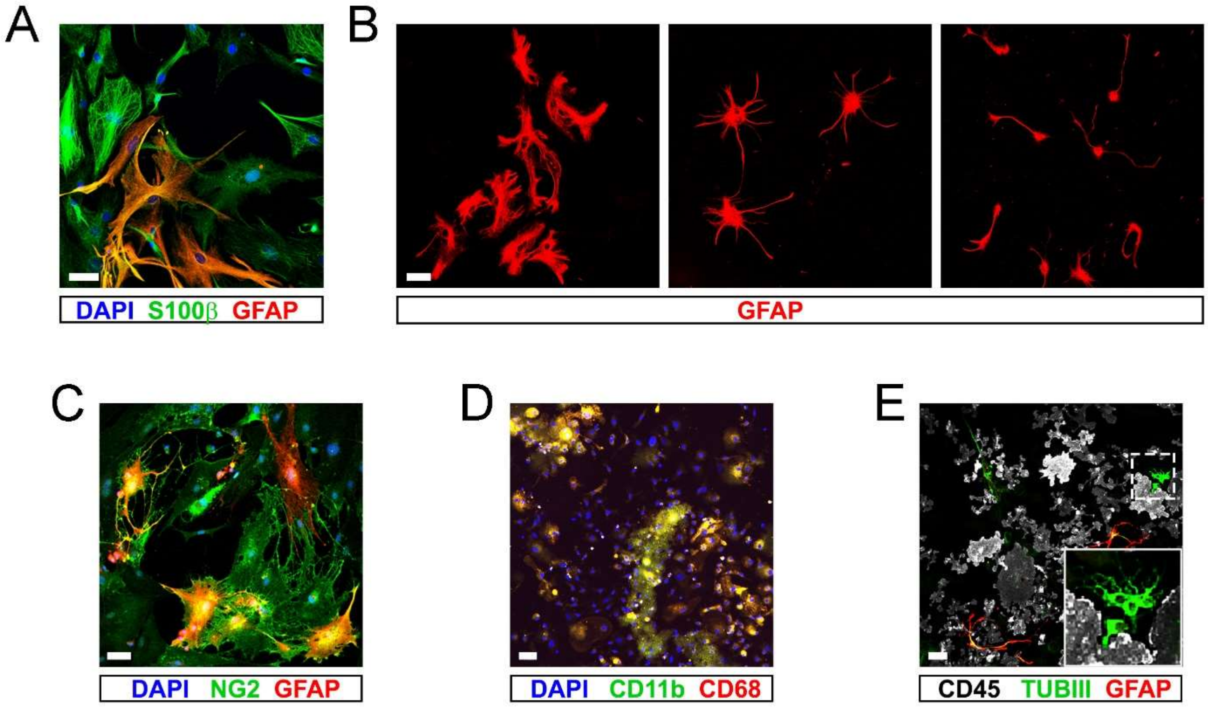


**Figure S3. Cultures contain diverse phenotypes of reactive glia and occasional neurons. (A)** Fluorescence microscopy image showing co-expression of S100β (green) and GFAP (red) in astrocytes from 3 DPI RCs at 5 DIV, with DAPI (blue) labeling nuclei. **(B)** Micrographs displaying the morphological diversity of $GFAP^+$ astrocytes in the same cultures. **(C)** Fluorescence microscopy image highlighting astroglial cells co-expressing GFAP (red) and NG2 (green), with arrows indicating double-positive cells. DAPI (blue) labels nuclei. **(D)** Myeloid cell populations in 1 DPI RCs at 5 DIV, visualized with CD11b (green) and CD68 (red) staining, alongside DAPI (blue) for nuclei. **(E)** Fluorescence microscopy image showing $TUBIII^+$ neurons (green) in in 3 DPI RCs at 8 DIV, along with $CD45^+$ immune cells (white) and $GFAP^+$ astrocytes (red). The inset provides a magnified view of $TUBIII^+$ neurons. Scale bars: 50 μm.

**A** Top 50 enriched proteins in RCs vs. HFs

| Order nº | Protein | Log2 ratios |
|---|---|---|
| 1 | Hbb-b1 | 8.64 |
| 2 | Serpina3n | 8.25 |
| 3 | Ctsd | 6.82 |
| 4 | Gap43 | 6.56 |
| 5 | Hba | 5.97 |
| 6 | Basp1 | 5.70 |
| 7 | Lgmn | 5.45 |
| 8 | Hexb | 5.32 |
| 9 | Mmp12 | 5.25 |
| 10 | Apoe | 5.04 |
| 11 | Cd5l | 4.72 |
| 12 | Ctsb | 4.39 |
| 13 | Lyz2 | 4.35 |
| 14 | Cox6c | 4.22 |
| 15 | Lpl | 4.21 |
| 16 | Ca2 | 4.09 |
| 17 | Got2 | 4.02 |
| 18 | Ptgr3 | 3.92 |
| 19 | Cep162 | 3.84 |
| 20 | Ctsl | 3.78 |
| 21 | Ctss | 3.75 |
| 22 | Igfbp6 | 3.70 |
| 23 | Stx1b | 3.68 |
| 24 | Rpl29 | 3.53 |
| 25 | Trem2 | 3.46 |
| 26 | Snca | 3.45 |
| 27 | Psap | 3.44 |
| 28 | Prdx1 | 3.44 |
| 29 | Col24a1 | 3.40 |
| 30 | Akap12 | 3.38 |
| 31 | Prrt2 | 3.30 |
| 32 | Brd9 | 3.27 |
| 33 | Dld | 3.04 |
| 34 | Hspb1 | 3.01 |
| 35 | Fsip2 | 2.95 |
| 36 | Fmod | 2.95 |
| 37 | Fuca1 | 2.93 |
| 38 | Fstl1 | 2.92 |
| 39 | Ppt1 | 2.89 |
| 40 | Got1 | 2.87 |
| 41 | Cct6b | 2.84 |
| 42 | Hyi | 2.82 |
| 43 | Lipa | 2.76 |
| 44 | Pkm | 2.76 |
| 45 | Loxl2 | 2.76 |
| 46 | Gsta4 | 2.74 |
| 47 | Ppp3r1 | 2.64 |
| 48 | H1-2 | 2.61 |
| 49 | Jpt1 | 2.58 |
| 50 | Ctsh | 2.58 |

**B** Common enriched proteins in RCs vs. P5As and RCs vs. HFs

| Order nº | Protein | Log2 ratios RCs vs. P5As | Log2 ratios RCs vs. HFs |
|---|---|---|---|
| 1 | Hbb-b1 | 8.27 | 8.64 |
| 2 | Hexb | 6.08 | 5.32 |
| 3 | Mmp12 | 5.99 | 5.25 |
| 4 | Ctsd | 5.90 | 6.82 |
| 5 | Basp1 | 5.67 | 5.70 |
| 6 | Hba | 5.29 | 5.97 |
| 7 | Lpl | 4.85 | 4.21 |
| 8 | Cd5l | 4.84 | 4.72 |
| 9 | Lgmn | 4.75 | 5.45 |
| 10 | Ctsb | 4.68 | 4.39 |
| 11 | Lyz2 | 4.32 | 4.35 |
| 12 | Got2 | 4.29 | 4.02 |
| 13 | Ctss | 4.27 | 3.75 |
| 14 | Col24a1 | 4.17 | 3.40 |
| 15 | Pkm | 4.05 | 2.76 |
| 16 | Stx1b | 3.87 | 3.68 |
| 17 | Fmod | 3.63 | 2.95 |
| 18 | Psap | 3.45 | 3.44 |
| 19 | Trem2 | 3.44 | 3.46 |
| 20 | Cox6c | 3.42 | 4.22 |
| 21 | Igfbp6 | 3.42 | 3.70 |
| 22 | Prdx1 | 3.34 | 3.44 |
| 23 | Ca2 | 3.31 | 4.09 |
| 24 | Gap43 | 3.16 | 6.56 |
| 25 | Ctsl | 3.15 | 3.78 |
| 26 | Loxl2 | 3.05 | 2.76 |
| 27 | Ptgr3 | 2.94 | 3.92 |
| 28 | Fstl1 | 2.91 | 2.92 |
| 29 | Ppp3r1 | 2.80 | 2.64 |
| 30 | Snca | 2.80 | 3.45 |
| 31 | H1-2 | 2.78 | 2.61 |
| 32 | Hspb1 | 2.78 | 3.01 |
| 33 | Ctsh | 2.60 | 2.58 |
| 34 | Prrt2 | 2.59 | 3.30 |
| 35 | Lipa | 2.29 | 8.25 |
| 36 | Serpina3n | 2.29 | 2.76 |

**C**

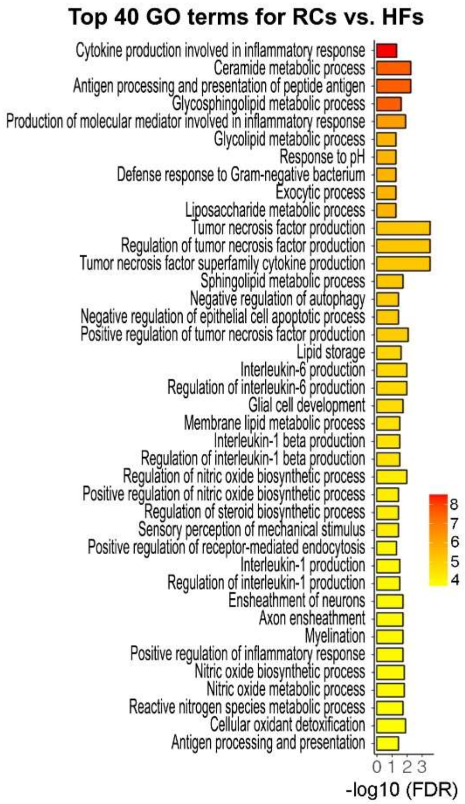


Extracellular matrix remodeling and structural integrity
Protein intracellular remodeling for tissue repair
Metabolism, oxidative stress and cell death
Cell proliferation and growth
Inflammation and immune response

**Figure S4. The extracellular medium from RCs shows a gliosis-like profile compared to both, HFs and P5As. (A)** Table showing the top 50 upregulated genes in RCs compared to HF. Genes associated with injury response are grouped and color-coded by function, as indicated in the legend. Note the red gradient which denotes levels of induction (standardized log2 ratios integrated by category; p ≤ 0.05, p-value). **(B)** Table showing the common genes upregulated in RCs compared to both P5As and HFs. **(C)** Table displaying the top 30 ranked GO terms for biological processes in RCs compared to HFs.

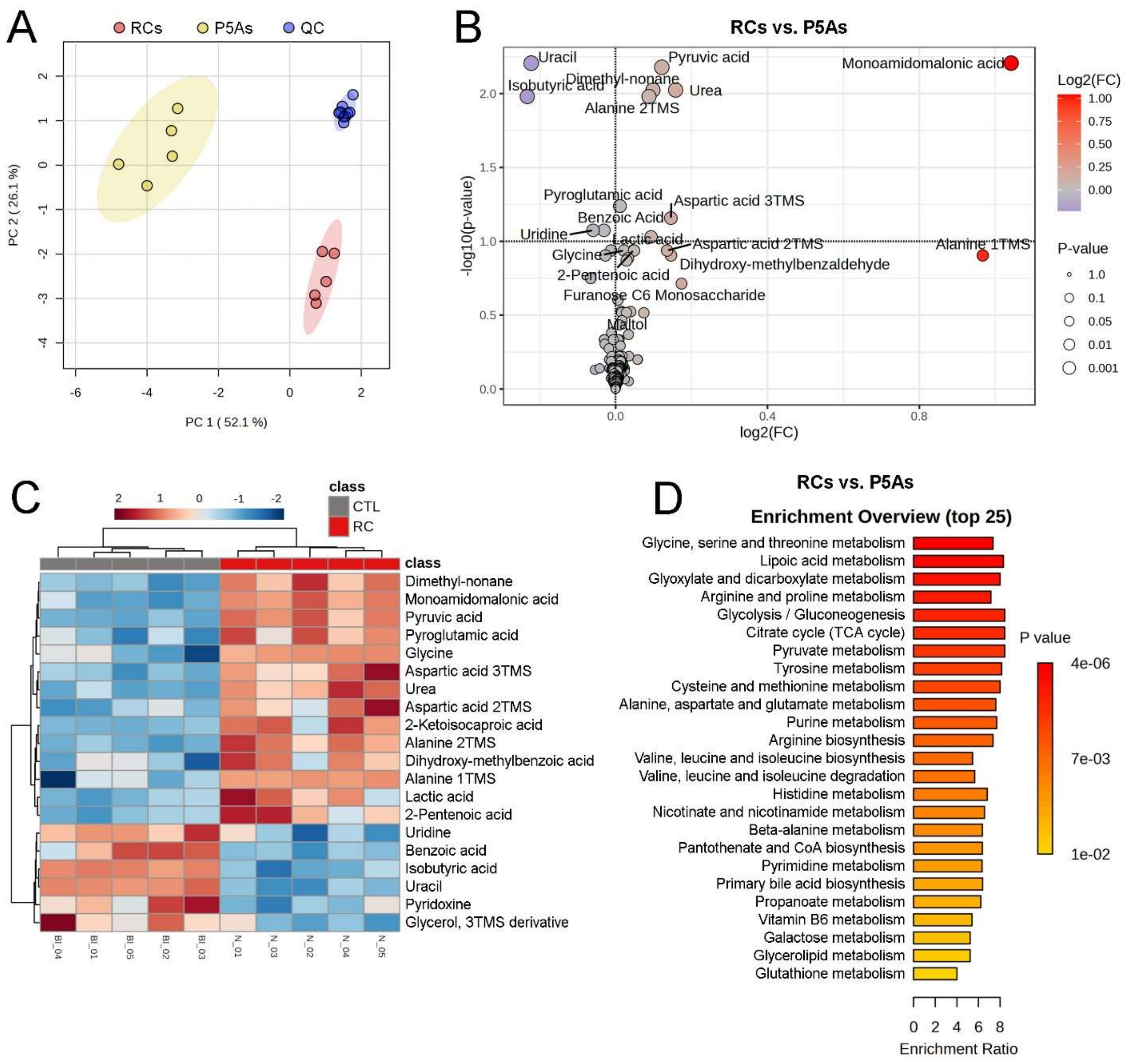


**Figure S5. Metabolomic analysis highlights injury-associated metabolic profiles distinguishing RCs from P5As. (A)** PCA plot illustrating the separation between the analyzed populations: P5As and RC. The distinct clustering of P5As and RC reflects divergent metabolic profiles, while the tightly clustered quality control (QC) samples (in blue) confirm consistent data quality throughout the experiment. **(B)** Volcano plot highlighting metabolites significantly upregulated or downregulated in RCs compared to P5As. Each point represents a metabolite, with color intensity denoting the fold change magnitude (log2(FC)) and point size representing the p-value significance. Key metabolites with significant changes are labeled. **(C)** Heatmap showing the relative abundance of 20 selected metabolites in RCs compared to P5As controls. Rows correspond to individual metabolites, and columns to samples. Metabolites are clustered based on abundance patterns, with a color gradient indicating z-scores: red for higher abundance and blue for lower abundance. **(D)** Pathway enrichment analysis of the top 25 enriched metabolic pathways in RC compared to control conditions. The enrichment ratio and p-value are indicated for each pathway, highlighting key metabolic processes associated with the RC medium. When multiple derivatives of the same metabolite are formed during the derivatization process, they are named according to the number of active oxygen atoms in the original molecule that have been replaced by trimethylsilyl groups (TMS). MA, monoamidomalonic acid; 2-KIC, 2-Ketoisocaproic acid; 2OH-methylbenzaldehyde, Dihydroxy-methylbenzaldehyde.

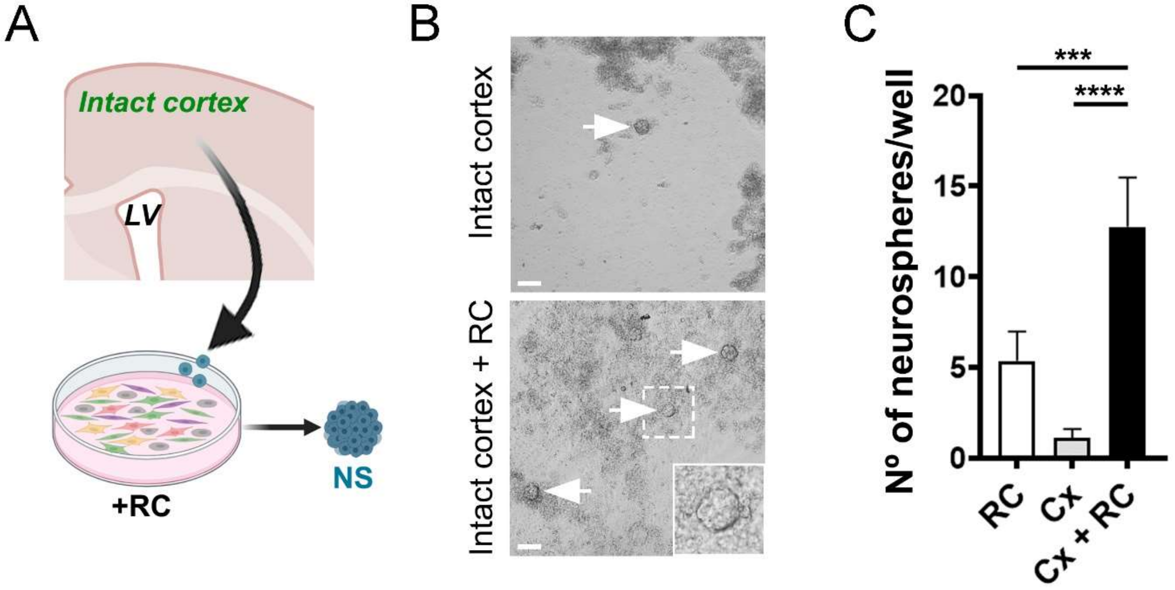


**Figure S6. Reactive cells induce intact-cortex cells to form neurospheres**. **(A)** Schematic of intact cortical cells cocultured with reactive cells under the neurosphere formation protocol. **(B)** Representative images of neurospheres formed by intact cortical cells cultured alone (upper panel) or cocultured with reactive cells (lower panel). **(C)** Quantification of neurospheres generated by reactive cells from injured cortex cultured alone, intact cortical cells cultured alone, and intact cortical cells cocultured with reactive cells. Statistical analysis: one-way ANOVA with FDR correction ** $p<0.01$; *** $p<0.001$.

**Supplementary Video 1 | Method for inducing controlled stab wound lesions in the cortex of adult mice.**

The video demonstrates the precise trajectory of the surgical lancet blade used to create a grid pattern of stab wound lesions in the visual cortex. Starting from the defined coordinates relative to Bregma (anteroposterior -1.6 mm to -2.5 mm; mediolateral -1.5 mm to -2.6 mm), the lancet is moved approximately 1 mm back and forth along both the anteroposterior and mediolateral axes to produce a total of 12 lesions, 6 in each direction. The diagram highlights the leading and trailing edges direction used for the cuts: in the anteroposterior direction, the blade moves parallel to its edge, creating precise stab wounds, while in the mediolateral direction, the blade moves perpendicularly to its edge, resulting in scratch-like lesions rather than deep cuts. This approach is suitable for inducing lesions in either unilateral or bilateral hemispheres (see Figure 1A).

**Supplementary Video 2 | Time-lapse imaging of neuronal conversion from oligodendroglial-lineage cells in reactive cultures.** The movie depicts a Sox10::GFP-positive cell (green), probably an OPC, transduced with Neurog2-RFP (red) undergoing neuronal reprogramming over 10 days, visualized across brightfield, GFP, and RFP channels as indicated. Yellow arrowheads depict the tracked cell across different channels as it transitions from a glial to a neuron-like phenotype, marked by neurite extension and gradual GFP signal attenuation, the latter indicative of Sox10 downregulation. The RFP zoom panel highlights detailed morphological changes of the tracked cell (labeled with a "1" in black), including the extension of neuronal processes. Scale bars, 150 μm.

**Supplementary Video 3 | Time-lapse imaging of astroglial-to-neuronal conversion in RCs.** Movie shows an astroglial cell marked by GFAP::GFP (green) transitioning towards a neuronal-like phenotype over 10 days, following Neurog2-RFP transduction (red). Yellow arrowheads depict the targeted cell losing GFP signal as GFAP expression diminishes, along with the development of extended processes characteristic of neurons. The RFP zoom panel highlights fine structural changes in the tracked cell (marked with a '2'), illustrating the acquisition of neuron-like morphology. Scale bars, 150 μm.
